# Dual blockade of LILRB1 and LILRB2 enhances antiviral immune responses in SIV infection

**DOI:** 10.64898/2026.03.03.709246

**Authors:** Florian Meurisse, Sixtine Coindre, Anne Wijkhuisen, Romain Marlin, Laure Fournier Le Ray, Juliette Pons, Laurent Abi-Rached, Anne-Sophie Gallouet, Francis Relouzat, Mael Gourves, Asier Saez-Cirion, Hisashi Arase, Gerard Zurawski, Sandra Zurawski, Nathalie Dereuddre-Bosquet, Roger Le Grand, Stéphanie Simon, Olivier Lambotte, Benoit Favier

## Abstract

Restoring effective antiviral immunity remains a major challenge in HIV infection. Among emerging immune checkpoint molecules, the inhibitory receptors LILRB1 and LILRB2 have been proposed as therapeutic targets, yet their in vivo function remains undefined due to the lack of cross-reactive blocking antibodies for relevant preclinical models. To address this, we developed a dual-specific blocking monoclonal antibody, mac20G10, targeting cynomolgus macaque LILRB1 and LILRB2 and assessed its immunomodulatory activity in an SIV model of infection.

Pharmacodynamics analyses demonstrated that mac20G10 persisted in circulation and engaged target myeloid cells for up to 14 days without detectable adverse effects. A single administration prior to SIVmac251 infection enhanced early myeloid immune activation, characterized by increased frequencies of CD80^+^ pDC and CD80^+^ monocyte/macrophage subsets in blood and lymphoid tissues. These changes were accompanied by increased plasma levels of IFN-λ, IL8, and IL-1RA during acute infection. Although viral replication remained unchanged, mac20G10 treatment promoted the development of SIV-specific memory CD8⁺ T-cell responses.

Together, these findings provide in vivo evidence that LILRB1 and LILRB2 function as myeloid immune checkpoints restraining antiviral priming, supporting this pathway as a rational target for combination immunotherapeutic strategies aimed at achieving durable HIV remission during analytic treatment interruption.

## Introduction

Following the success of immunotherapy in cancer, strategies based on blocking immune checkpoints with monoclonal antibodies (mAb) are emerging as a promising treatment in HIV infection ^1–5^. LILRB1 and LILRB2 are inhibitory receptors mainly expressed by monocytes, macrophages and dendritic cells (DC) that plays an important role in the regulation of immune responses modulating the progression of infectious diseases and cancer ^6,7^. These receptors interact with classical and non-classical MHC class I (MHC-I) molecules, delivering inhibitory feedback signals that restrain myeloid cell activation ^6–12^. Importantly, these LILRB1/B2-mediated immune inhibitions can be reversed by blocking mAb in vitro ^13,14^. However, investigation of their functions has been limited in vivo due to an absence of clear orthologs in murine models ^7,15^.

HIV and SIV infections are characterized by an early dysregulation of myeloid immune responses that contributes to suboptimal adaptive response and disease progression ^16–19^. We and others have previously demonstrated that expression of LILRB2 and their MHC-I ligands is enhanced on myeloid cell subsets in early HIV and SIV infection ^20,21^. This enhancement of the LILRB2/MHC-I inhibitory axis directly impairs the shaping of immune responses against HIV in vitro ^22–24^. Furthermore, studies in people living with HIV exhibiting distinct progression profiles have shown that the strength of the LILRB2/MHC-I interaction positively correlates with DC dysfunction and accelerated disease progression ^25^. Elevated levels of soluble MHC-I molecules in the plasma of people living with HIV further amplify this inhibitory pathway by engaging LILRB2 on monocytes and DC, thereby attenuating CD80 and CD86 up-regulation and resulting in impairment of T cell priming ^26,27^. Although the contribution of LILRB1 in HIV infection remains less well defined, accumulating evidence suggests that multiple pathogens, including dengue virus, HCMV and *Plasmodium falciparum*, exploit LILRB1 engagement to attenuate immune responses and promote persistence ^28–30^. These convergent aspects underscore the potential of targeting the LILRB1/MHC-I interaction to reinvigorate antiviral immunity in chronic infections.

Despite compelling in vitro evidence supporting LILRB2 and LILRB1 as therapeutic targets in HIV infection, their roles remain to be characterized in vivo. To address this gap, the cynomolgus macaque (*Macaca fascicularis*) SIVmac251 infection model closely mirrors key features of human HIV infection, including the rapid peak in plasma viral load (around day 14 post-infection) followed by stabilization during the chronic phase. This well-established model recapitulates major aspects of HIV pathogenesis in humans and provides a robust platform for investigating disease mechanisms in vivo. Notably, it has been instrumental in characterizing the dysregulation of myeloid immune cell subsets during early SIV infection and their downstream impact on adaptive immune responses ^16,17,21,31–35^.

Here, we report the development of a monoclonal antibody targeting cynomolgus macaque LILRB1 and LILRB2, designed to block their interaction with MHC-I molecules. A single administration of the LILRB1/B2 blocker prior to SIV challenge enhanced early myeloid immune activation and elicited a specific cytokine signature, which was associated with improved induction of SIV-specific CD8⁺ T cell responses during the chronic phase of infection. Together, these findings highlight the therapeutic potential of targeting LILRB1 and LILRB2 in HIV infection.

## Results

### Development of a blocking monoclonal antibody targeting cynomolgus macaque LILRB1 and LILRB2

An anti-LILRB1/B2 mAb was generated using cynomolgus macaque LILRB1 as immunogen (**Fig. 1A**). To this end, a soluble cynomolgus macaque LILRB1 extracellular region fused to the human IgG1 Fc portion was used to immunize mice. Hybridoma supernatants were initially screened for recognition of cynomolgus macaque LILRB1 (mafa-LILRB1) and LILRB2 (mafa-LILRB2) by ELISA and flow cytometry. Monoclonal antibodies of interest were then isolated and assessed for their specificity toward LILRB1 and LILRB2, as well as their ability to block interactions between LILRB1 or LILRB2 and Mafa-A1*063 SIV^gag^ tetramers by flow cytometry. Clone 20G10 was selected based on its superior combined specificity for LILRB1/B2 and blocking activity. Flow cytometry analysis demonstrated that clone 20G10 recognizes mafa-LILRB1 and mafa-LILRB2, but not LILRB3 or LILRB4, nor LILRA1, LILRA2, or LILRA4 (**Fig. 1B**). In addition, the 20G10 mAb efficiently blocked mafa-LILRB2 binding to Mafa-A1*063 SIV^gag^ tetramers (**Fig. 1C**).

**Figure 1.**
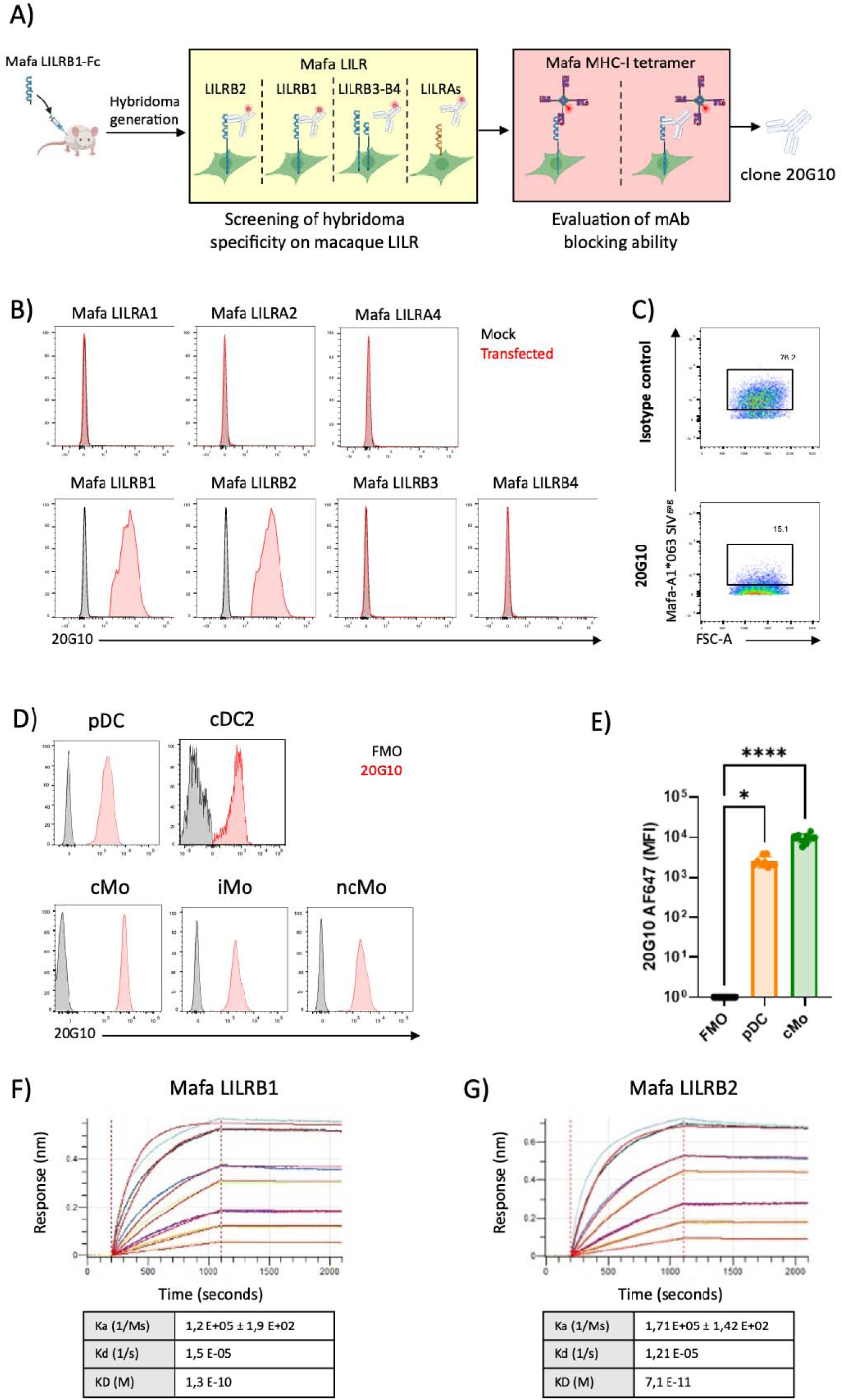
**Development of an anti-LILRB1 and LILRB2 dual blocker mAb (clone 20G10) specific of cynomolgus macaque.** A) Schematic overview of the strategy used to generate a dual-blocking anti-LILRB1/LILRB2 monoclonal antibody. B) Determination of 20G10 mAb specificity using HEK293T expressing cynomolgus macaque receptor LILRA1, LILRA2, LILRA4, LILRB1, LILRB2, LILRB3 or LILRB4. Non-transfected control cells are represented in grey. C) 20G10 mAb blocking ability was assessed using a system of interaction between cynomolgus macaque LILRB2 expressed by HEK293T and cynomolgus macaque MHC-I tetramers (Mafa-A1*063) harboring a SIV^gag^ peptide. Stained cells were analyzed by flow cytometry. A significant reduction of LILRB2 binding to Mafa-A1*063 tetramer demonstrated the blocking activity of the antibody. D) Cell distribution of 20G10 mAb staining was assessed using flow cytometry. PBMCs from cynomolgus macaque were used to characterize 20G10 staining of pDC, cDC2 and monocyte subsets. cMo = classical monocytes, iMo = inflammatory monocytes and ncMo = non-classical monocytes. 20G10 staining are represented in red and controls in grey. E) Histograms representing MFI of pDC and cMo (n=5) following staining with 20G10-AF647, comparisons with negative controls were carried-out using Dunn’s statistical test; * = p<0.05, **** = p<0.0001. F-G) Binding parameters of 20G10 mAb to cynomolgus macaque LILRB1-Fc (F) and LILRB2-Fc (G) were assessed using Bio-layer Interferometry. Sensograms and fitted curves are displayed as well as the summary of the kinetic parameters. Cynomolgus macaque LILRB1-Fc and LILRB2-Fc were tested at seven concentrations ranging from 50 to 0.78nM in twofold serial dilutions, represented by different colours in the sensogram.

Consistent with previous human studies, staining of cynomolgus macaque PBMCs with fluorescent 20G10 mAb confirmed that LILRB1 and LILRB2 are predominantly expressed by plasmacytoid dendritic cells (pDC), cDC2, and monocytes subsets (**Fig. 1D-E and Fig. S1**). The 20G10 mAb also cross-reacted with monocytes isolated in rhesus macaque (*Macaca mulatta*), another pre-clinical model widely used to investigate HIV pathogenesis (**Fig. S2**).

The affinities of the 20G10 mAb for cynomolgus macaque LILRB1-Fc and LILRB2-Fc were measured by interferometry. High affinity was observed for both inhibitory receptors (K_D_= 1.3×10^-10^ M and K_D_=7.1×10^−11^ M, respectively) (**Fig. 1F-G**).

Overall, these data indicate that 20G10 specifically recognizes and blocks cynomolgus macaque LILRB1/B2 on myeloid cells in vitro, highlighting its potential as an immune-therapeutic candidate.

### Humanization and optimization of the 20G10 mAb for in vivo use

Since mouse mAb can be immunogenic in primates, the mAb clone 20G10 was modified by replacing mouse constant fragments with human IgG1 (ch20G10). Mutations LALA and YTE were incorporated to attenuate potential adverse events and increase circulating half-life in vivo ^36^ (**Fig. 2A**). Next, variable regions of ch20G10 were humanized to a degree of 87.8% (hu20G10). Finally, human Fc IgG1 regions were replaced by cynomolgus macaque constant Fc IgG1 regions (including the LALA and YTE mutations), in order to increase homology with cynomolgus macaque physiology (mac20G10). The effects of these modifications on K_D_ were assessed for cynomolgus macaque LILRB1 by interferometry. Affinity was not altered by the combined series of modifications, with optimized mac20G10 possessing a similar K_D_ to parental 20G10 isoform (**Fig. 2B**). Apparent K_D_ for cynomolgus macaque LILRB1 and LILRB2 were also measured by flow cytometry (**Fig. 2C-D**). Slight decreases of affinity of mac20G10 for cynomolgus macaque LILRB1 or LILRB2 was observed, compared to ch20G10 (0.5913 µg/mL to 2.075 µg/mL and 0.8779 µg/mL to 2.699 µg/mL, respectively).

**Figure 2.**
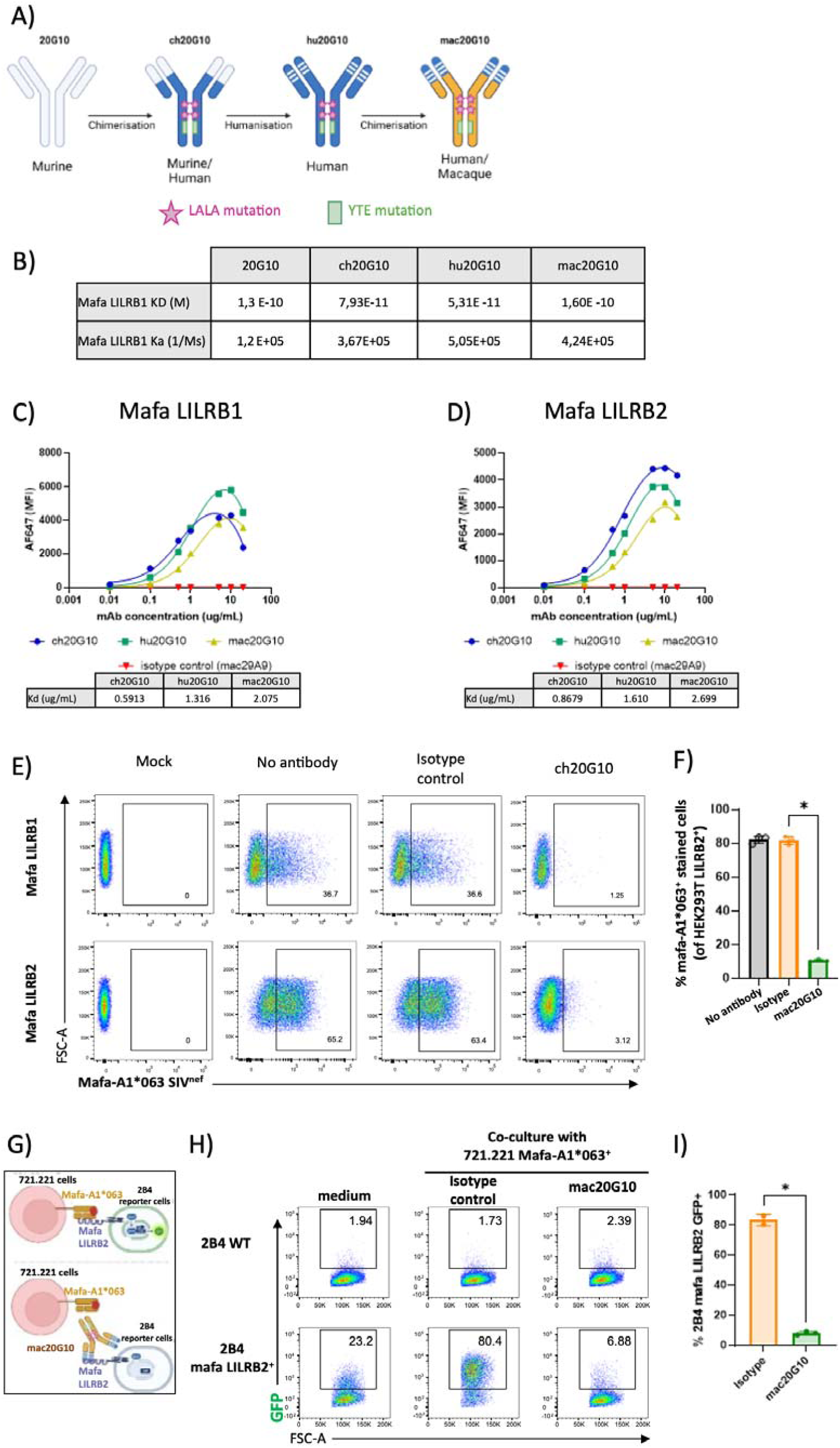
**Optimization of anti-LILRB1/B2 blocking antibody for in vivo use.** A) Schematic representation of the strategy used to optimize 20G10 mAb for cynomolgus macaque studies. The LALA mutation reduces the risks of adverse events by minimizing Fc receptor binding, while the YTE mutation enhances the antibody’s half-life in vivo by increasing affinity for FcRn. B) Summary of KD and Ka of 20G10, ch20G10, hu20G10 and mac20G10 for cynomolgus macaque LILRB1-Fc. KD were assessed using Bio-layer Interferometry. C-D) Apparent dissociation constants (Kd) of the different antibody variants determined by flow cytometry. HEK293T expressing either Mafa-LILRB1 or Mafa-LILRB2 were incubated with various concentrations of ch20G10, hu20G10 or mac20G10 to evaluate binding affinities to cynomolgus macaque LILRB1 (C) or LILRB2 (D). The mAb mac29A9 was used as control. E) ch20G10 mAb blocking ability was assessed using a system of interaction between cynomolgus macaque LILRB1 or LILRB2 expressed by HEK293T and cynomolgus macaque MHC-I tetramers (Mafa-A1*063) harboring a SIV^nef^ peptide. Stained cells were analyzed by flow cytometry. A significant reduction in tetramer binding demonstrates the efficacy of the antibody. F) Histograms showing the frequency of HEK293T LILRB2^+^ cells stained with Mafa-A1*063^+^ tetramers following the blocking assay performed in the absence of antibody (n=3) or in the presence of either an isotype control (n=3) or mac20G10 (n=3) mAb. Comparisons with the isotype control were carried-out using Dunn’s statistical test; * = p<0.05. G) Schematic representation of the coculture reporter cell system assay. H) mac20G10 mAb blocks the interaction of LILRB2 and MHC-I in coculture reporter cell system. The GFP signal is representative of a productive interaction between expressed receptor and a ligand. I) Histograms representing frequency of 2B4 mafa LILRB2^+^ GFP^+^ post coculture assay with medium alone (n=2) or in presence of 721.221 expressing Mafa-A1*063 and isotype control (n=2) or blocking antibody (n=3). Comparisons with isotype controls were carried-out using Dunn’s statistical test; * = p<0.05.

Blocking assays demonstrated efficient ch20G10-mediated inhibition of LILRB1 or LILRB2 binding to the Mafa-A1*063 SIV^gag^ tetramer (36.6% vs 1.25% for LILRB1 and 63.4% vs 3.12% for LILRB2; isotype control vs ch20G10) (**Fig. 2E**). This blocking capacity was preserved following antibody optimization (medians at 83.1% and 82.2% vs 10.2%; without mAb and isotype vs mac20G10, p<0.05) (**Fig. 2F**). The blocking activity of mac20G10 was further assessed using a coculture assay between NFAT-GFP reporter cells expressing LILRB2, which produce GFP upon recognition of Mafa-A1*063 presented by 721.221 cells (**Fig. 2G**). Coculture resulted in robust GFP expression, reflecting productive receptor-ligand interaction, and this signal was effectively abrogated by preinbucation with mac20G10 but not with the isotype control. Importantly, mac20G10 displayed no agonistic activity, as no GFP expression was detected in reporter cells cultured with mac20G10 (**Fig. 2H, 2I**).

Overall, these data indicate that the optimized mac20G10 is endowed with high affinity and blocking properties well suited for testing therapeutic applications.

### Blocking LILRB1/B2 mAb binds to target myeloid immune cells without inducing adverse events in cynomolgus macaque

Either ch20G10 or mac20G10 was administered to groups of healthy cynomolgus macaques, and pharmacodynamic parameters were characterized (**Fig. 3A**). The persistence of circulating mAb in plasma evaluated by ELISA show that ch20G10 and mac20G10 concentrations remained elevated for at least 14 days (**Fig. 3B**). In parallel, the interaction between mAb and myeloid targets was monitored using a flow cytometry-based competition assay, as previously described ^37^ (**Fig. 3C-D**). Both ch20G10 and mac20G10 remained bound to LILRB1/B2 on pDC for at least 14 days (**Fig. 3E-F**). Accordingly, a strong correlation was observed between plasma persistence of the mAb and pDC binding (r = - 0.7408, p <0.0001) (**Fig. S3A**). Similarly, anti-LILRB1/B2 mAb remained bound to the surface of cDC2 and monocytes subsets for at least 14 days (**Fig. 3G and 3H**). Furthermore, competition assays performed on pDC and cDC2 subsets isolated from axillary lymph node (LN) biopsies of *n = 2* cynomolgus macaques treated with ch20G10 seven days earlier demonstrated that the mAb remained bound to both cell population within the LN at day 7 post-injection (**Fig. 3I**).

**Figure 3.**
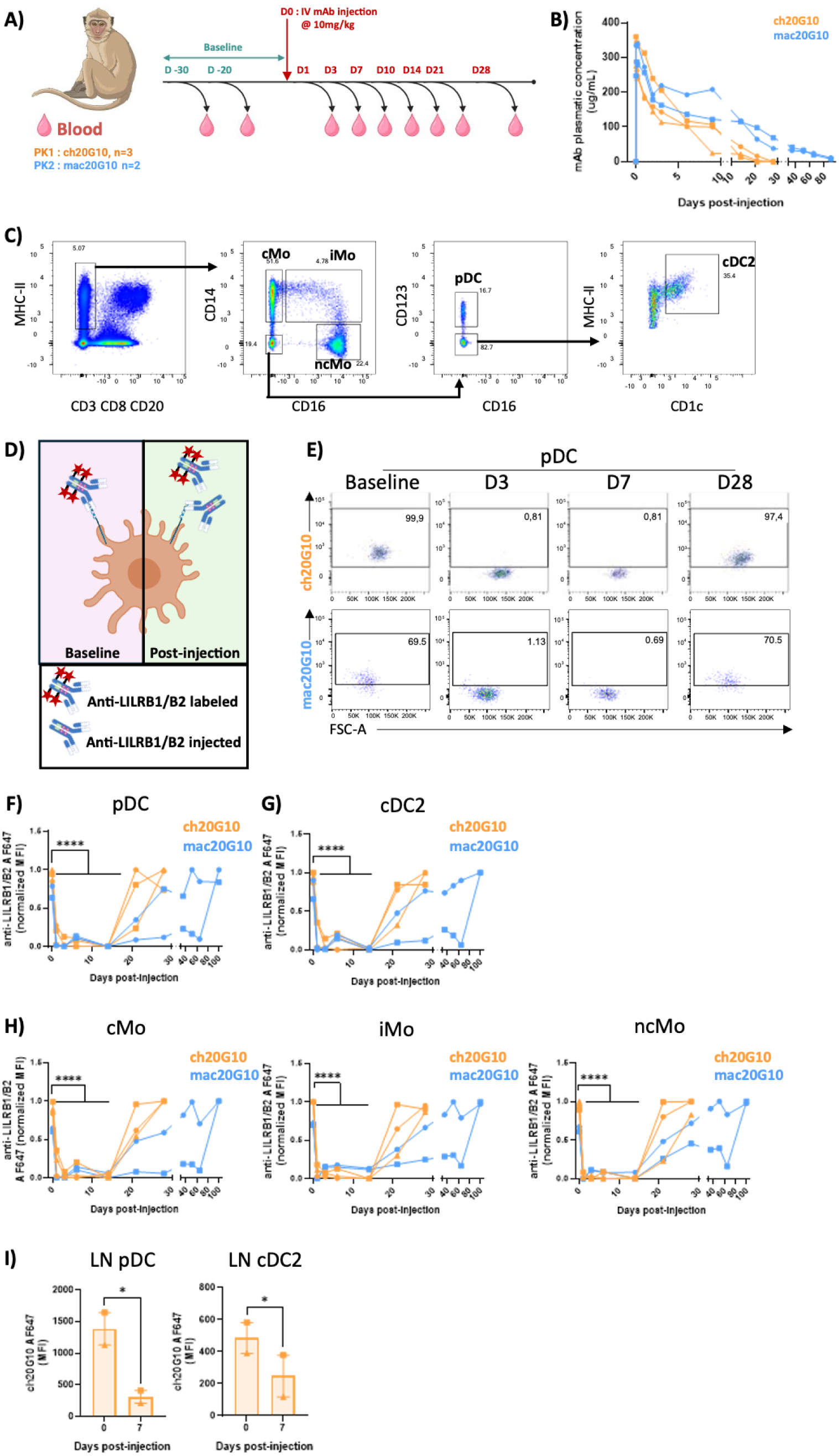
**Pharmacodynamics of ch20G10 and mac20G10 in cynomolgus macaques.** A) Schematic representation of the pharmacodynamic studies schedule. Two group were constituted. The first group received mAb ch20G10 (n=3, 10 mg/kg, orange) and the second group received mAb mac20G10 (n=2, 10 mg/kg, blue). B) Plasma levels of ch20G10 (orange) and mac20G10 (blue) were monitored throughout the studies by ELISA. C) Flow cytometry strategy used to monitor mAb binding to cMo = classical monocytes, iMo = inflammatory monocytes and ncMo = non-classical monocytes, pDCs, and cDC2 during the pharmacodynamic studies. D) Schematic representation of ex vivo competition assay performed by flow cytometry to follow antibody persistence at the surface of myeloid cell subsets. E) Representative flow cytometry dot plots of competition staining experiments of pDC during the follow-up of the animals at baseline, D3, D7 and D28 post-administration of ch20G10 or mac20G10. F-H) Surface persistence of mAb on pDC, cDC2 and monocytes subsets was monitored by competition staining assay. Values were normalized and compared to the baseline using mixed-effect analysis, followed by post-hoc analysis; **** = p<0.0001. I) Competition staining assays carried-out by flow cytometry with ch20G10 on pDC and cDC2 from axillary lymph nodes (n=2 individuals). Conditions were compared using a Mann Whitney statistical test; * = p<0.05.

Toxicity analysis in cynomolgus macaques treated with ch20G10 or mac20G10 indicated that administration of either antibody had no impact on physiological or biochemical parameters. Furthermore, no depletion of the myeloid or lymphoid hematopoietic cell compartments was observed during the study (**Fig. S3 B-D**).

Taken together, these data demonstrate that the recombinant ch20G10 and the optimized mac20G10 antibodies bind to LILRB1/B2 on myeloid cell subsets for at least two weeks following administration to cynomolgus macaques, without any detectable adverse events. These findings validate mac20G10 as a powerful tool for investigating the role of LILRB1/B2 in cynomolgus macaque preclinical models of human infectious diseases.

### Effect of mac20G10 treatment during SIVmac251 infection

To ensure optimal tissue distribution and engagement of target immune cells prior to infection, cynomolgus macaques received a single dose of either anti-LILRB1/B2 blocking antibody mac20G10 or the isotype control mac29A9, two days before SIVmac251 challenge. Immune responses and viral load dynamics were subsequently monitored in these animals (**Fig. 4A**). No differences of plasma viral load evolution or CD4^+^ T cell dynamics were observed between the two groups (**Fig. 4B and S4**). However, mac20G10 remained detectable by ELISA in the plasma of treated animals for 10 to 14 days after SIV infection (corresponding to 12 to 16 days after mAb administration) (**Fig. 4C**). The flow cytometry strategy enabled the simultaneous characterization of pDC, cDC1, cDC2 and monocytes subsets in peripheral blood (**Fig. S5A**). Concordantly with anti-LILRB1/B2 kinetics measured in plasma (**Fig. 4C**), competition assays indicated that mac20G10 remained bound to circulating pDC, cDC2 and classical monocytes (cMo) for 10 to 14 days post-infection (**Fig. 4D**). Similar results were observed for pDC and CD163^+^CD14^+^ macrophages isolated from peripheral LN using dedicated flow cytometry strategy (**Fig. S5B**), with binding detected on day 7 post-infection (**Fig. 4E**).

**Figure 4.**
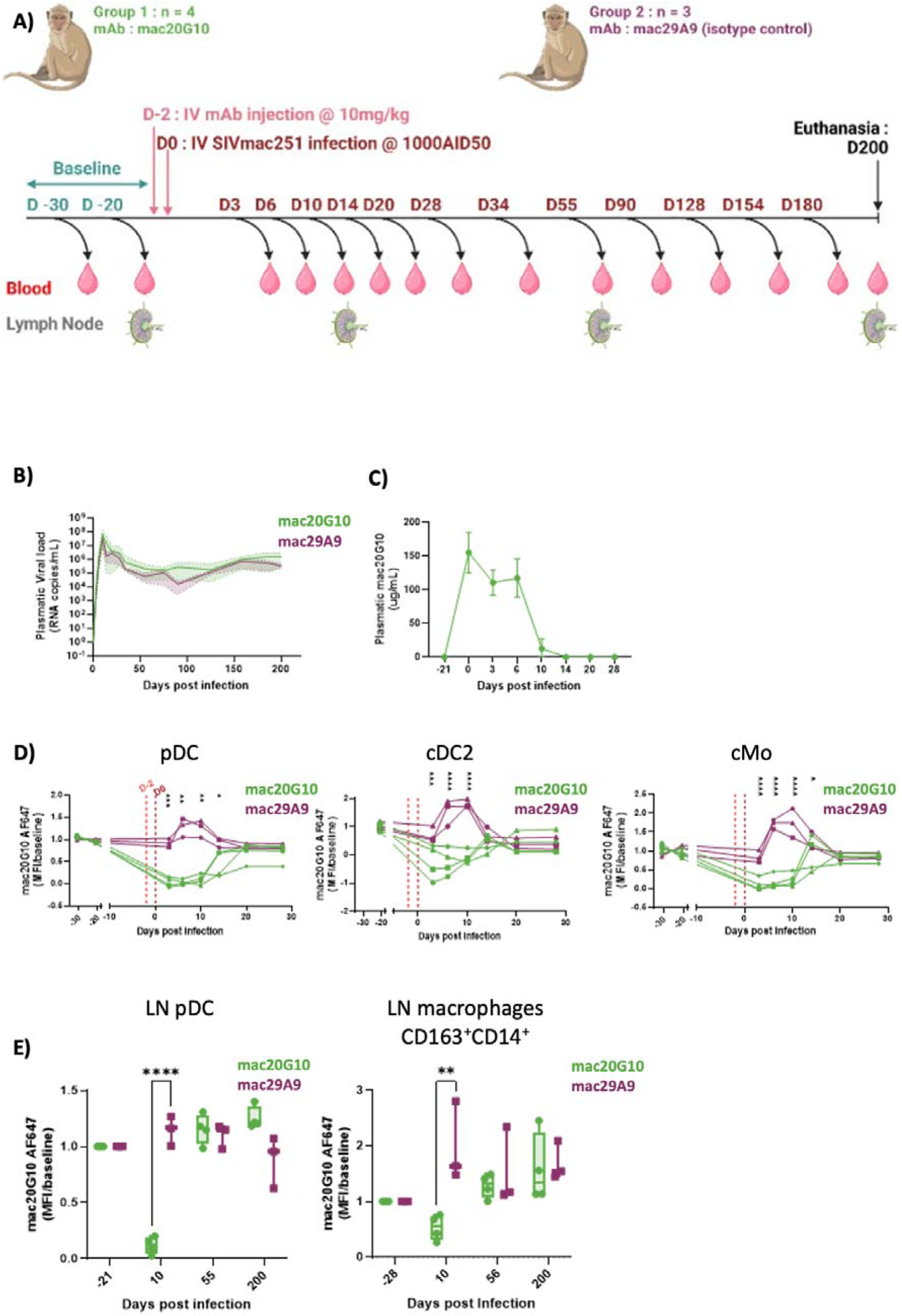
**Longitudinal analysis of viral load and mac20G10 pharmacodynamics in SIV-infected cynomolgus macaques.** A) Schematic representation of the study aiming to evaluate the impact of mac20G10 on immune responses in SIV model. Two groups of cynomolgus macaques were used. One received the blocking anti-LILRB1/B2 mac20G10 (n=4, green) while the second received the isotype control mac29A9 (n=3, purple) both before infection. Animals were followed for 200 days after infection. B) Follow-up of SIV plasma viral load post infection in both groups by RT-qPCR. C) Follow-up of mac20G10 persistence in plasma by ELISA. D) Surface persistence of mac20G10 on pDC, cDC2 and classical monocytes (cMo) monitored by competition staining assay. Normalized values are reported relative to baseline. The two groups were compared using a Kruskal-Wallis test, followed by post-hoc analysis; * = p<0.05, ** = p<0.01; *** = p<0.001; **** = p<0.0001. E) Characterization of mac20G10 binding to pDC and macrophages (CD163^+^ CD14^+^) in lymph nodes, by competition staining assay. Normalized values are reported relative to baseline. Data were compared using a Kruskal-Wallis test, followed by post-hoc analysis; ** = p<0.01, **** = p<0.0001.

Blood cell count did not differ between control and mac20G10-treated animals (**Fig. S6A**). Consistent with previous finding, MHC-I ligands were upregulated on cDC2 and monocyte subsets during early SIV infection (**Fig. S6B**) ^21^.

These data indicate that a single administration of mac20G10 is sufficient to efficiently bind to LILRB1/B2^+^ myeloid subsets during at least 10 days post SIV-infection, but has no effect on evolution of viral infection, nor on circulating immune cell type composition.

### mac20G10 promotes CD80 expression on pDCs and monocytes/macrophages and increases plasma IFN-**λ**, IL8 and IL-1RA in early SIV infection

To further evaluate the impact of LILRB1/B2 blockade on myeloid cell subsets during early cynomolgus macaque SIV infection, expression of key costimulatory molecules CD80, CD86 and CD40 were assessed on myeloid populations from blood and LN by flow cytometry. Treatment with mac20G10 enhanced the frequency of CD80^+^ pDC at day 6 (medians: 16.3% vs 4.7%; mac20G10 vs control, p<0.01) and day 10 (medians: 58.6% vs 45.9%; mac20G10 vs control, p<0.01) post-infection (**Fig. 5A-B**). In addition, when compared to baseline, the proportion of CD80^+^ pDC and cDC2 were significantly increased at day 6 post-infection in the mac20G10-treated group (pDC medians: mac20G10-treated 0.5% to 16.3% p<0.01, vs control 0.2% to 4.7%; cDC2 medians: mac20G10-treated 9.1% to 21.4% p<0.05, vs control 6.0% to 12.4%) (**Fig. 5B-C**). However, at day 10 post-infection for cDC2 no significant differences were observed between the two groups (medians: 56.9% vs 41.7%; mac20G10 vs control, p=0.08).

**Figure 5.**
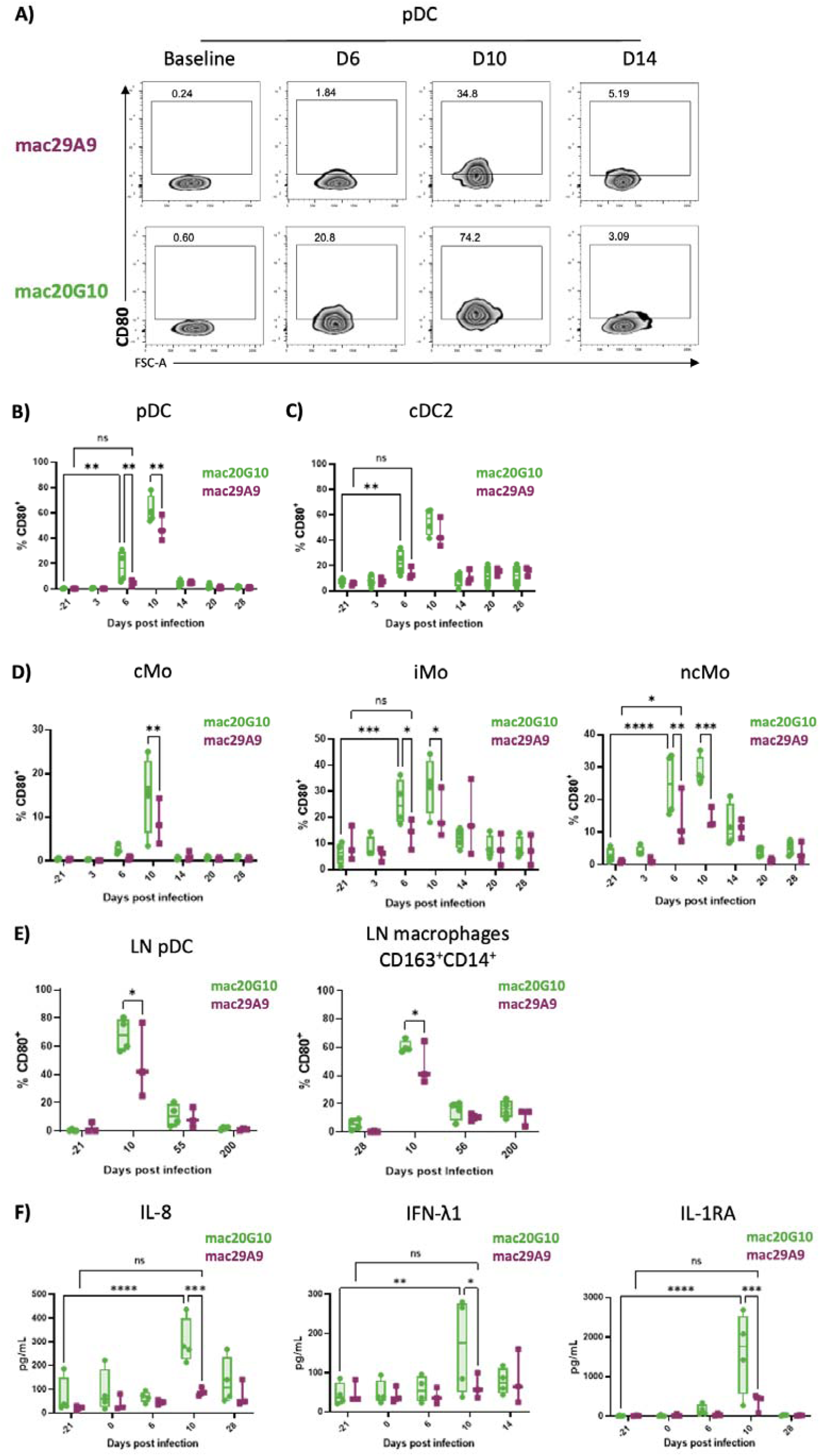
**Follow-up of the impact of mac20G10 on CD80^+^ pDC, cDC2 and monocytes/macrophages frequencies and on plasma cytokine production during early SIV infection.** A) Representative zebra plots of flow cytometry follow-up of CD80^+^ pDC subset during early SIV infection in cynomolgus macaques treated with mac20G10 or isotype control. B) Follow-up by flow cytometry of CD80^+^ pDC population during early SIV infection in both groups. Groups and time points were compared using a Kruskal-Wallis test, followed by post-hoc analysis; ** = p<0.01. C) Follow-up of the CD80^+^ cDC2 population during early SIV infection by flow cytometry. Groups and time points were compared using a Kruskal-Wallis test, followed by post-hoc analysis; ** = p<0.01. D) Follow-up of CD80^+^ monocytes sub-populations during early SIV infection by flow cytometry. cMo = classical monocytes, iMo = inflammatory monocytes and ncMo = non-classical monocytes. Groups and time-points were compared using a Kruskal-Wallis test, followed by post-hoc analysis; * = p<0.05, ** = p<0.01, *** = p<0.001, **** = p<0.0001. E) Follow-up by flow cytometry of CD80^+^ pDC and macrophages (CD163^+^ CD14^+^) populations in lymph node during SIV infection. Groups were compared using a Kruskal-Wallis test, followed by Dunn’s statistical test; * = p<0.05. F) Follow-up of plasma concentrations of IL8, IFN-λ and IL-1RA during early SIV infection in cynomolgus macaques treated with mac20G10 or isotype control. Time points and groups were compared using a Kruskal-Wallis test, followed by post-hoc analysis; * = p<0.05, ** = p<0.01, *** = p<0.001, **** = p<0.0001. The group treated with mac20G10 is represented in green (n=4) and mac29A9 control isotype group in purple (n=3).

The frequency of CD80^+^ cMo subsets were higher at day 10 post SIV infection for the mac20G10-treated group (cMo medians: 15.8% vs 8.2%, iMo medians: 32.6% vs 17.8% and ncMo medians: 27.3% vs 12.6%; mac20G10 vs control, p <0.05) (**Fig. 5D**). As observed for pDC, the proportion of CD80^+^ intermediary (iMo) and non-classical (ncMo) monocytes had already increased by day 6 post-infection after mac20G10 treatment.

Higher frequencies of CD80^+^ pDCs and macrophages were elicited in mac20G10-treated animals at day 10 post SIV-infection (pDC medians: 67.9% vs 41.7%, and macrophages medians: 58.9% vs 40.8%; mac20G10 vs control, p<0.05) (**Fig. 5E**).

No significant impact of mac20G10 treatment was observed via analysis of CD40 and CD86 expression on pDC, cDC2 or monocyte subsets (**Fig. S7**). In addition, mac20G10 treatment did not seem to affect cDC1 cell count, MHC-I and costimulatory molecule expression during early SIV infection (**Fig. S8**).

Plasma cytokines concentrations were also analyzed during SIV infection. For isotype-treated SIV-infected animals, the cytokine expression profile was similar to the profiles previously observed in untreated SIV infection ^38^. However, production of IL8, IFN-λ and IL-1RA were enhanced at day 10 post infection animals treated with mac20G10 (IFN-λ medians: 175.3 pg/mL vs 56.6 pg/mL, p<0.05; IL-8 medians: 274.1 pg/mL vs 85.5 pg/mL and IL-1RA medians: 1754.6 pg/mL vs 445.3 pg/mL, p<0.001; mac20G10 vs control) (**Fig. 5F**). No difference was observed for other soluble immune mediators between the two groups (**Fig. S9 and S10).**

Taken together, these results indicate that a single administration of mac20G10 induces a higher frequency of CD80^+^ myeloid cells in blood and LN, which is accompanied by concomitant increases in a specific set of cytokines during early SIV infection.

### mac20G10 administration enhances the development of SIV^gag^-specific CD8^+^ memory T cell responses

To assess the impact of mac20G10 treatment on CD8^+^ T cell activation and IFN-γ production during SIV infection, we stimulated cynomolgus macaque PBMCs ex vivo with a pool of SIV^gag^ peptides in baseline and 180 days post-infection. The level of CD8^+^ T cell activation was then assessed by measuring the expression of the CD69 marker or the intracellular production of IFN-γ by flow cytometry. Following stimulation, a higher frequency of CD69^+^ CD8^+^ T cells was observed in the chronic phase in the group that received mac20G10 (medians: 0.2% to 4.1% for mac20G10, p<0.05; 0.2% to 2.07% for control, p>0.05; baseline to chronic phase) (**Fig. 6A**). Similarly, an enhancement of IFN-γ production was observed in the mac20G10 treated group (medians: 0% to 1.2% for mac20G10, p<0.05 and 0% to 0.51% for control, p>0.05; baseline to chronic phase) (**Fig. 6A**).

**Figure 6.**
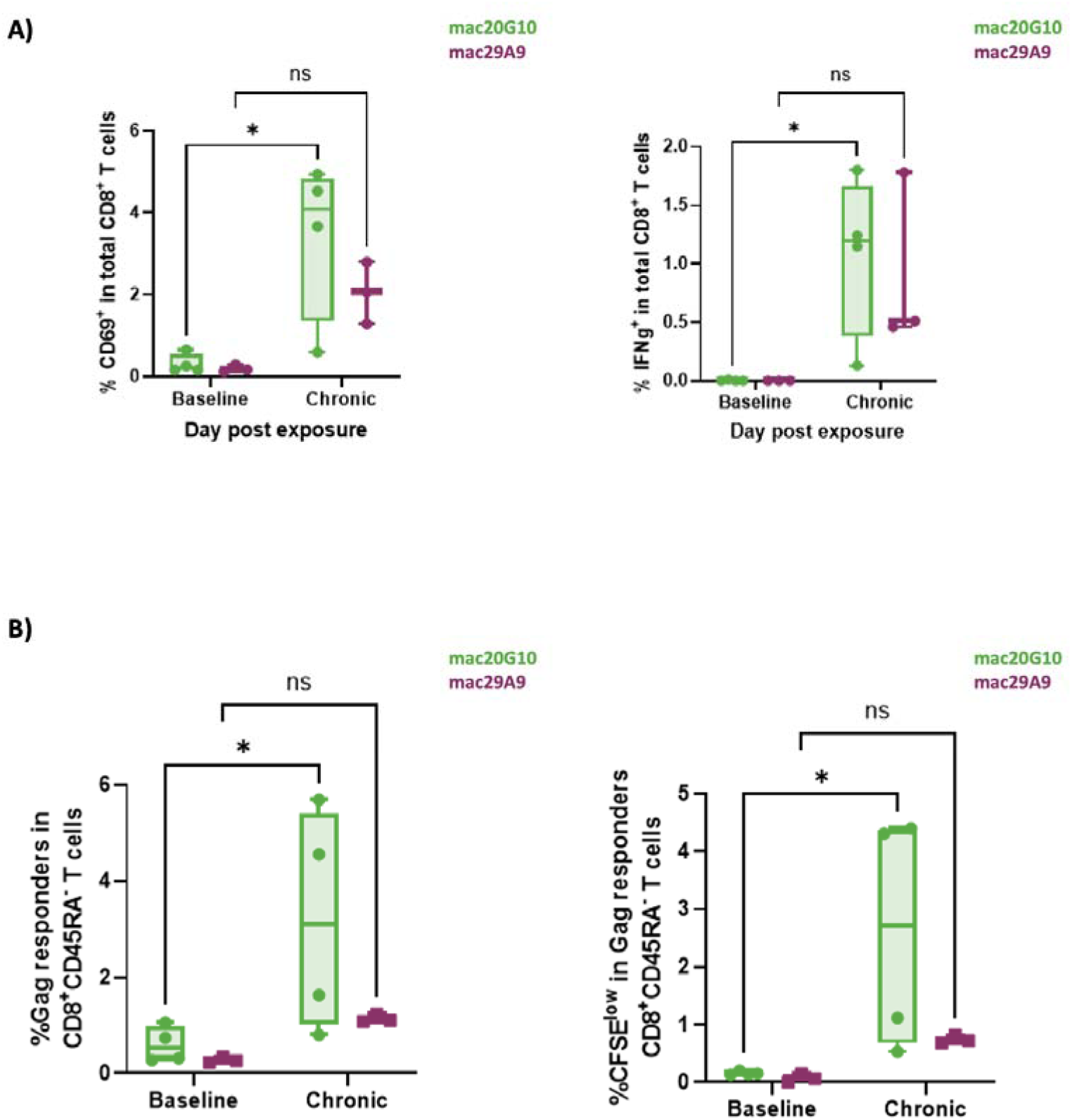
**Evaluation of mac20G10 treatment effects on CD8^+^ T-cell antiviral responses during the chronic phase of SIV infection.** A) Evaluation of specific CD8^+^ T cell responses to SIV^gag^ peptides ex vivo stimulation by measuring CD69 activation marker expression (left) and intracellular IFN-γ production (right) by flow cytometry. Time points and groups were compared using a Kruskal-Wallis test, followed by post-hoc analysis; * = p<0.05, ** = p<0.01, *** = p<0.001. B) Ex vivo evaluation of SIV specific memory CD8 T cell enrichment and ability to proliferate over 6 days following SIV peptides stimulation. Time points and groups were compared using a Kruskal-Wallis test, followed by post-hoc analysis; * = p<0.05 mac20G10 treated group is represented in green (n=4) and control isotype group in purple (n=3).

Further analysis of the memory potential of CD8^+^ T cells, demonstrated robust generation of SIV-specific CD8^+^ T memory cells in the group treated with mac20G10, which was not significant in isotype-treated animals (medians: 0.5% to 3.1% for mac20G10, p<0.05, and 0.3% to 1.1% for control, p>0.05; baseline to chronic phase). Furthermore, SIV-specific CD45RA^-^ CD8^+^ T cells from mac20G10-treated animals were characterized by cytokine production following expansion upon SIV-peptide stimulation in contrast to the control group (medians: 0.1% to 2.7% for mac20G10, p<0.05, and 0.1% to 0.7% for control, p >0.05; baseline to chronic phase) (**Fig. 6B and Fig. S11**).

These results demonstrate that a single administration of mac20G10, just prior to SIV infection, elicits enhanced CD8^+^ T cell memory responses against SIV during the course of SIV infection.

## Discussion

Immune checkpoint blockade has revolutionized cancer immunotherapy but remains underexplored in chronic viral infections, where efforts have largely focused on reversing T cell exhaustion ^39,40^.

Here, we provide in vivo evidence that blockade of the myeloid inhibitory receptors LILRB1 and LILRB2 enhances antiviral immune responses during SIV infection, identifying a myeloid immune checkpoint axis that operates upstream of adaptive immunity and shapes durable antiviral responses. The engagement of LILRB1 and LILRB2 has been associated with dysfunctional immunes responses and disease progression in HIV/SIV infection but also in other infectious diseases, yet in vivo functional evidence has been limited by the absence of suitable preclinical tools ^6^. By developing a dual LILRB1/B2-blocking antibody suitable for cynomolgus macaques, we were able to directly assess the role of these receptors in shaping immune responses in a physiologically relevant model of HIV infection.

Previous reports highlighted how alterations in costimulatory molecule expression and enhanced LILRB2/MHC-I inhibitory signaling were hallmarks of DC and monocyte/macrophage dysregulation in SIV and HIV infection that could lead to degraded immune responses ^16,20–27,41^. Analysis of the lymphoid tissues from people living with HIV and SIV infected macaques also underlined defective expression of costimulatory molecules by DC and up-regulation of LILRB2 ^21,42,43^. Our data indicate that LILRB1/B2 blockade enhances CD80 expression on pDC, cDC2 and macrophages in lymph nodes during early SIV infection, further supporting a role for the LILRB1/B2 inhibitory pathway in the regulation of myeloid cell activation. In contrast, no changes in CD86 and CD40 expression by myeloid cell subsets were observed. Such discrepancies between costimulatory molecules was previously reported in cDC2 and macrophages during early SIV infection, reflecting the selective regulation of costimulatory gene expression by the LILRB1/B2 pathway ^16^.

Enhanced myeloid activation was accompanied by increased production of IL8, IL-1RA and IFN-λ during early infection, indicating that disruption of LILRB1/B2 - MHC-I interactions augments functional cytokine responses in this context. As these cytokines are primarily produced by myeloid cells, these data support a model in which LILRB1/B2 blockade reshapes the early myeloid immune response, thereby improving conditions for effective immune priming. Because monocytes/macrophages and DC are the main immune cells producing IL8, IL-1RA and IFN-λ ^44–46^, future single-cell level studies will be important to determine whether CD80^+^ myeloid subsets also exhibit improved functional capacities, including enhanced ability for cytokine production.

We previously reported positive correlations between CD80 expression on DC and the level of CD8^+^ T cells activation in human elite HIV controllers ^35^. Our data herein further revealed that inhibition of LILRB1/B2 signaling enhances the development of SIV-specific CD8^+^ T cell responses, thus providing a proof-of-concept that improving early myeloid responses benefits adaptive immune responses in SIV infection. The enhancement of the SIV-specific CD8^+^ T cell response we observed, which endured into the chronic phase, suggests that LILRB1/B2 blockade is an important strategy to produce durable antiviral immunity during immune response priming.

Despite these immunological effects, plasma viral load was not significantly altered. This outcome is likely explained by the experimental context, which employed a high-dose intravenous challenge resulting in rapid viral replication that exceeds the capacity of early immune modulation to constrain viremia. Under such conditions, enhancement of immune responses does not necessarily translate into acute virological control, as previously reported for T cell-targeted therapy immune checkpoint blockade in SIV infection, highlighting the distinction between immunological enhancement and direct virological control ^37,47,48^. In addition, HIV/SIV rapidly establishes viral reservoirs, often before full maturation of innate and adaptive immunity, which may limit the capacity of a single intervention to reduce early viremia ^49,50^.

Rather than limiting translational relevance, these findings refine the optimal positioning of LILRB1/B2 blockade. By enhancing memory CD8_⁺_ T cell responses, this strategy may be particularly effective in settings of partial viral suppression, such as chronic infection under antiretroviral therapy, where immune-mediated control of viral rebound is critical. Accordingly, LILRB1/B2 blockade may be best suited for use in analytical treatment interruption (ATI) protocols, either alone or in combination with T cell–directed checkpoint inhibitors ^51,52^.

To date, immune checkpoint blockade in the SIV macaque model included anti-CTLA-4 and anti-PD1 ^37,47,53–57^. Administration of blocking anti-CTLA-4 prior to SIV infection lead to increased viral replication in mucosal sites ^54^. This effect prompted the discontinuation of further studies using anti-CTLA-4. Anti-PD1 was administrated either in the early or chronic phase with up to six multiple injections, with or without antiretroviral therapy ^47,55–57^. This treatment improved CD8^+^ T cell responses, which were characterized by improved IFN-γ production after ex vivo stimulation with SIV^gag^ peptides ex-vivo. The extent of CD8^+^ T cell enhancement induced by sequential anti-PD1 treatment was close to that observed in our study after a single administration of LILRB1/B2 blocker. In light of these results, multiple injections of anti-LILRB1/B2 may improve the efficiency of immunomodulation, thereby promoting even better adaptive immune response generation. However, such a protocol could also elicit anti-drug antibodies (ADA) and would require pre-evaluation of mac20G10 immunogenicity elicited by repeat administrations prior to conducting such a study ^58^.

Combinatorial checkpoint blockade strategies are increasingly explored as a means to restore immune competence and achieve functional viral control during ATI ^59–61^. The data presented here support the rationale for combining LILRB1/B2 blockade with PD-1 inhibition to simultaneously reinvigorate myeloid function and reverse T cell exhaustion in ATI. Such synergy could enhance both the quality and durability of antiviral immune responses. Supporting this rationale, recent studies demonstrated that SIV-infected macaques under ART exhibited significant viral load reduction when treated with combination of anti-PD1 and anti-IL10 inhibitory mAb administered as a multiple-injection regimen, just before and after ART interruption ^62^.

Beyond HIV infection, LILRB1 and LILRB2 can regulate immune responses that shape the outcome of various infectious and malignant diseases. That absence of detectable adverse effects following LILRB1/B2 blockade, together with sustained antibody persistence on target cells in the blood and lymphoid organs, highlights the translational potential of this strategy. The dual-blocking monoclonal antibody developed in this study provides a valuable tool for interrogating LILRB1/B2 biology and evaluating therapeutic interventions in macaque models of human disease.

In summary, our study demonstrates that dual blockade of the myeloid inhibitory receptors LILRB1 and LILRB2 enhances early immune activation and establishes durable antiviral CD8_⁺_ T cell responses during SIV infection. Although insufficient to achieve viral control alone, these findings provide compelling in vivo evidence that myeloid immune checkpoint blockade can potentiate antiviral priming and support its integration into combinatorial strategies aimed at achieving sustained, immune-mediated viral control in HIV infection, particularly in the context of analytic treatment interruption.

## Material and Methods

### Ethics statements

Cynomolgus and rhesus macaques were imported from Mauritius and housed in facilities at the Infectious Diseases Models and Innovative Therapies (IDMIT) center (CEA site in Fontenay-aux-Roses, France). Non Human primate studies were approved by the ethics committee “Comité d’éthique en expérimentation animale” No. 44, and by the “*Ministère de l’Education Nationale, de l’Enseignement Supérieur et de la Recherche*” (France) (*Reference :* APAFIS#20525-2019050616506478 and APAFIS #39765-2022121215145877 v1). All animal treatments and procedures were performed according to French national regulations and, by association, European Union regulations under the direct supervision of national veterinary inspectors (*CEA accreditation No. D92-032-02*) (*European Directive 2010/63, recommendation No. 9*), in compliance with the Standards for Human Care and Use of Laboratory Animal Welfare (*OLAW, USA; under OLAW Assurance No. A5826-01* and #F22-00556).

### Mice

Biozzi mice were bred at the animal care unit of the CEA (Gif sur Yvette, France). All experiments were performed in compliance with French and European regulations on the care of laboratory animals (European Community Directive 86/609, French Law 2001-486, 6 June 2001) and with the agreements of the Ethics Committee of the Commissariat à l’Energie Atomique (CEtEA ‘Comité d’Ethique en Expérimentation Animale’ n° 44) no. 12-026 and 15-055 delivered to S. S. by the French Veterinary Services and CEA agreement D-91-272-106 from the Veterinary Inspection Department of Essonne (France).

### Human samples

Blood samples of healthy subjects were obtained from anonymous donors via the “*Etablissement Français du Sang*” (EFS).

### Cell lines, cell culture and transduction

Human 721.221 MHC-I deficient cell line and murine 2B4 reporter cells were cultured at 37°C and 5% CO2 in RPMI 1640 GlutaMAX (Gibco) containing 10% fetal bovine serum (FBS) (Sigma-Aldrich) and 1% Penicillin-Streptomycin (Gibco). Human 293T embryonic kidney (HEK293T) cells were cultured at 37°C and 5% CO2 in DMEM GlutaMAX (Gibco) complemented with 10% FBS and 1% Penicillin-Streptomycin (Gibco). Cells were periodically tested for mycoplasma contamination using MycoAlert Mycoplasma Detection kit (Lonza).

HEK293T, 721.221 and 2B4 cell lines expressing either LILR or MHC-I of interest were obtained through lentiviral transduction. Briefly, cells were concentrated around 5×10^6^ cells/mL and lentiviral particles were added with a MOI of 15. Suspensions were incubated at 37°C for 1 hour within gentle mix each 15min. At the end, medium was added in order to dilute cells at 0.25×10^6^ cells/mL. After 16 hours of incubation, medium was changed to fresh medium and cells were plated in T25. After 4 days of culture, cells were stained with either anti-HLA-I (clone W6/32) or anti-LILR in PBS + 0.5% BSA for 30min at 4°C. Then, cells were washed, and sorted for the expression of LILRs or MHC-I and GFP reporter gene using ARIA Fusion 2 cell Sorter (BD).

### Cynomolgus macaque LILR-Fc fusion protein production

The LILR-Fc fusion proteins were produced using LILRB1 cynomolgus macaque cDNA sequence (Reference XP_045236898.1). The cDNAs encoding the extracellular domain of the macaque LILRB1 or LILRB2 were fused to a hIgG1-Fc by molecular biology. The plasmid constructions were transfected into ExpiHEK293T cells and supernatants were harvested to purify LILRB1-Fc or LILRB2-Fc proteins by affinity chromatography.

### LILRB1/B2 monoclonal antibody generation, optimization and production

Monoclonal antibodies (mAbs) were raised in Biozzi mice by immunizing with cynomolgus macaque LILRB1-Fc (30 μg per injection) fusion protein. Mice eliciting the highest anti-antibody response against LILRB1 were given an intravenous boost injection 3 days before being sacrificed for splenic B cell fusion, according to Köhler and Milstein ^63^. Hybridoma culture supernatants were screened for antibody production, specificity for cynomolgus macaque LILRB1 and LILRB2, blocking capacity, by enzyme immunoassay, flow cytometry, or both methods. Selected hybridomas were subsequently cloned by limiting dilution. Monoclonal antibodies were produced in hybridoma supernatants and further purified using AKTA protein purifier. Purity of monoclonal antibodies was assessed by SDS-PAGE in reducing and non-reducing conditions using an Agilent 2100 Bioanalyser. Endotoxin levels in purified mAbs were determined by Pierce LAL Chromogenic Endotoxin Quantitation Kit (Thermo Fisher Scientific) according to the manufacturer’s protocol.

cDNA corresponding to variable regions of selected hybridoma clone 20G10 were amplified by RT-PCR and sequenced by the Sanger method. ch20G10 was obtained by grafting cDNA sequences of variable regions into a vector containing human IgG1 Fc constant regions harboring LALA (L234A, L235A) and YTE (M252Y, S254T, T256E) mutations.

The humanization of 20G10 variable regions was performed by deep mutation scanning (DMS) and yeast surface display (YSD) as described previously ^64^. Humanized 20G10 variants were produced and tested in vitro before selection of the best candidate.

The mac20G10 was obtained by grafting selected the humanized 20G10 sequence to cynomolgus macaque IgG1 constant chain sequences including LALA and YTE mutations. The isotype control mac29A9 was generated using variable region sequences specific to a hapten.

Monoclonal antibodies, clone ch20G10 and mac20G10, were produced at large-scale in ExpiCHO cells. Then mAb were purified from supernatant using affinity chromatography with protein A in low endotoxin conditions. Quality controls were assessed using SDS-PAGE, spectrophotometry and SEC-HPLC. The protein sequences of the antibody variants are included in a related patent application (EP 3 981 789).

### Immunoenzymatic screening of anti-LILRB1 and -LILRB2 hybridoma supernatants

Enzyme immunoassays (EIA) were performed by transferring 50 μl of diluted hybridoma culture supernatants into 96-well, flat-bottom microplates (MaxiSorp^TM^ immunoplate, Nunc) coated with goat anti-mouse immunoglobulin antibody (Jackson ImmunoReseach), and incubated overnight at 4 °C. After washes, biotinylated mLILRB1-Fc or mLILRB2-Fc fusion proteins (50 ng/ml) were added (100 μl per well) and plates were reacted for 1 h at room temperature (RT). Plates were washed and reacted for 30 min at RT with 100 μl per well of 1 EU/ml of acetylcholinesterase (AChE)-labeled streptavidin. After several washes, AChE activity was revealed by Ellman’s colorimetric method ^65^ by measuring absorbances at 414 nm after 1 h. All reagents were diluted in EIA buffer (0.1 M phosphate buffer [pH 7.4] containing 0.15 M NaCl, 0.1% BSA, and 0.01% sodium azide). Plates coated with proteins were saturated in EIA buffer (18 hours at 4°C) and washed with washing buffer (0.1 M potassium phosphate [pH 7.4] containing 0.05% Tween 20).

### Affinity determination of mAbs

The affinities of mAbs for the different macaque LILRs-Fc fusion proteins were determined by Bio-layer Interferometry using the ForteBio system (Pall Laboratory). mAbs prepared at 10µg/ml in EIA buffer + 0.02% Tween 20 (Sigma-Aldrich) were dispensed in 96-well microplate. In another wells, LILR-Fc fusion proteins were each dispensed at 8 titrated concentrations. A glycine (Sigma-Aldrich, pH [1.4]) regeneration solution and EIA buffer + 0.02% Tween 20 for baseline stabilization and neutralization was also prepared. The plate was agitated at 1000 rpm over the entire course of the experiment. Prior to the binding measurements, the anti-mouse Fc (AMC) sensors tips were hydrated in EIA buffer + 0.02% tween 20. The sensor tips were then transferred to the EIA buffer + 0.02% Tween 20 for the baseline, after mAb – containing wells for the 300 sec loading step. After the baseline step in EIA buffer + 0.02% Tween 20 for 60 sec, the binding kinetics were measured by dipping the mAb-coated sensors into the wells containing LILR-Fc fusion protein at varying concentrations. The binding interactions were monitoring over 900 sec associated period and followed by a 900 sec dissociation period in the wells containing EIA buffer +0.02 % Tween 20. The AMC sensors tips were regenerated with wells containing glycine [pH1.4] and neutralized in the EIA buffer + 0.02% tween 20 between each binding cycle. The equilibrium dissociation constant (KD) was calculated using the ratio between the dissociation rate constant (koff) and the association rate constant (kon), obtained with global Langmuir 1:1 fit (Octet Data Analysis software, vHT.10).

Apparent Kd was also determined using flow cytometry. Shortly, HEK293T cells expressing macaque LILRB1 or LILRB2 were plated at 10^5^ cells per well in 96 U-wells plate and incubated with either anti-LILRB1/B2 mAb, isotype control, or without antibody for 1.5 h at 4°C, within periodic agitation. Then, cells were washed twice in PBS 0.5% BSA and stained with polyclonal anti-human IgG-AF647 antibodies (Jackson Immunology) for 30 min. Finally, cells were centrifuged at 200g for 7min and resuspended in Fixation buffer (Biolegend) diluted at 1:4 in 1X PBS and then analyzed by flow cytometry.

### Cross-reactivity assay

HEK293T cells expressing macaques LILR were used to assess cross-reactivity of anti-LILRB1/B2 mAb. The different macaque LILRs contained a DYKDDDDK Flag in N-terminal to check their expression at cell surface by flow cytometry with an anti-Flag APC (Miltenyi). Cells were incubated with anti-LILRB1/B2 mAb, or isotype control for 30 min at 4°C. Then, cells were centrifuged at 200g for 7 min and resuspended in Fixation buffer (Biolegend) diluted at 1:4 in 1X PBS.

### Blocking assay

HEK293T cells expressing macaque LILRB1 or LILRB2 were plated at 10^5^ cells per well in 96 wells plate and washed once. Cells were first incubated for 20min at RT with, either, anti-LILRB1/B2 mAb, isotype control or alone. Then, fluorescent tetramers Mafa-A1*063-PE carrying SIV^gag^ peptide (Clinisciences) were added for 30min at 4°C. Blocking ability was determined by flow cytometry, based on PE signal detection.

### Reporter assay

721.221-Mafa-A1*063 cells were co-incubated with either 2B4 reporter cells expressing macaque LILRB2 fusion protein or 2B4 parental cells at 1:1 ratio in 96 U-Wells plate. Then, cells were incubated at 37°C and 5% CO2 for 24 hours in presence of anti-LILRB1/B2 (20 ug/ml) or isotype control (20 ug/ml). At the end, cells were washed and stained with anti-human HLA-I (W6/32, Biolegend) and anti-human CD45 (HI30, BD) for 30min at 4°C to identify each cell lines and GFP production was quantified by flow cytometry.

### Pharmacodynamic studies

Two pharmacodynamic studies were carried-out with either ch20G10 (n=3) or mac20G10 (n=2) versions of anti-LILRB1/B2 blocking mAb. Antibodies were infused intravenously at 10mg/kg body weight at day 0. Blood samples were collected before mAb administration and up to 4 months following injection.

### Antibody administration and SIV infection

Seven female macaques were included in the study. They were 3 years old at the inclusion. The first group (n=4) and second group (n=3) were infused with either mac20G10 or isotype mac29A9 isotype control, respectively. Antibodies were administrated intravenously at 10mg/kg of body weight 48 hours before infection. Both groups were infected intravenously with 1,000 animal infectious dose 50% (1000AID_50_) of the isolate SIVmac251 on day 0. Macaques were periodically sampled, then euthanized 6 months after infection. Animals were negative for the MHC-I H6 haplotype, in order to prevent natural control SIVmac251 viral infection.

### Samples collection and processing

Blood for complete blood cell count, phenotyping and plasma assays was collected in EDTA tubes (Vacutainer, BD, USA). Blood for functional assays was collected in Cell Preparation Tube (CPT) heparin lithium tubes (Vacutainer, BD, USA). Blood for serum collection was collected in dry tubes (Vacutainer, BD, USA). Peripheral lymph nodes (LN) were harvested from the right and left inguinal and axillary nodes in clockwise direction. All procedures were performed under general anesthesia by intramuscular injection of ketamine (Imalgene 1000, 10 mg/kg) / medetomidine (Domitor, 0.5 mg/kg).

EDTA and dry tubes were centrifuged at 2100g for 10min, at room temperature, to obtain, respectively, plasma and serum. Plasma was used to follow circulating antibody concentration by a in-house ELISA, cytokines concentration by Luminex or Legendplex and viral load by RT-qPCR. Buffy coat was diluted at 1:1 within PBS 1X (Gibco). After sampling 100uL for whole blood staining, the cell mixture was completed with homemade hypotonic red blood cells lysis buffer at a ratio of 1:15. Whole blood leukocytes obtained were recovered by centrifugation at 200g for 10mins and cells were resuspended in PBS 1X + 0.5% BSA (Sigma-Aldrich). Cell count was assessed using a Vi-Cell XR automated hemocytometer (Beckman Coulter).

LN samples were divided in half with a scalpel before mechanical dissociation of single cells, using gentle pressure applied with a sterile syringe plunger on a 70 µm nylon filter. The filter was periodically washed with fresh RPMI 1640 GlutaMAX containing 10% FBS and 1% Penicillin-Streptomycin. Finally, LN cells were recovered by centrifugation at 200g (4°C) for 10 mins. Final cells suspensions were resuspended in PBS 1X + 0.5% BSA.

### Competition assay

Whole blood cells were stained with anti-LILRB1/B2 mAb labeled with AF647 (Antibody Labeling Kits Alexa Fluor 647, Thermo Fischer) for 30min at 4°C. Cells were analyzed using flow cytometry after washes and fixation. The absence of signal post-injection indicates that macaque LILRB1/B2 receptors are saturated by the antibody administered.

### Anti-LILRB1/B2 mAb distribution and reactivity cross-macaque

Staining of immune cell subsets by 20G10 from cynomolgus or rhesus macaques were performed by incubating whole blood cells with appropriate antibody cocktails for 30min at 4°C. Cells were analyzed using flow cytometry after washes and fixation. Antibodies are described in **Table S1**.

### Immune cells phenotyping and functional assays

Antibody cocktails were prepared in PBS 1X + 0.5% BSA, using previously calculated saturating concentrations. For each sample, whole blood cells were labeled with the appropriate antibody cocktail. For competition assay anti-LILRB1/B2 mAb labeled with AF647 was included. Whole blood cells were pre-incubated with the corresponding antibody cocktail for 30min at 4°C. Subsequently, whole blood staining tubes were lysed using Lysis/Fixation buffer (Biolegend) for 7 min at RT and washed with PBS 1X + 0.5% BSA and pelleted at 200g for 7 min. Cells were then fixed with Fixation buffer (Biolegend) diluted at 1:4 in PBS 1X. Antibodies are described in **Table S1**.

For functional assays, 1×10^6^ PBMCs were incubated overnight in presence of overlapping SIV^gag^ or SIV^nef^ peptides at a final concentration of 2ug/mL, at 37°C and 5% of CO^2^. Brefeldin A was added after 2 hours (final concentration: 10ug/mL) to inhibit cytokine secretion. At the end of the incubation, cells were washed, fixed, permeabilized (Cytofix/Cytoperm, BD) and stained. T cell responses were characterized by measuring the frequency of CD8 T cells expressing IFN-γ (clone B27, BD), TNF-α (clone Mab11, Biolegend), CD69 (clone FN50, Biolegend). The CD3 (clone SP34-2, BD), CD4 (clone L200, Becton Dickinson) and CD8 (clone BW135/80, Miltenyi) antibodies were used as lineage markers. For the assessment of memory potential, PBMCs were labelled with CFSE at 1 µM (Invitrogen, ref C34554)) and stimulated with a pool of optimal SIV peptides (2 µg/mL) for 6 days as previously described^66^. Twelve hours before completing the culture PBMCs were re-stimulated with the peptide pool. Brefeldin A (10 µg/mL; Invitrogen, ref 00-4506-51), Monensin (1 µg/mL; BD Biosciences, ref 554724) were added at this time.

Stained-cell suspensions were acquired with a 5 LASERs ZE5 cytometer (Biorad). FlowJo v10.10.0. was used for the analysis of all cytometry data.

### Analysis of immune soluble mediators

Cytokines and chemokines concentrations were quantified in EDTA plasma using Non-Human Primate Cytokine/Chemokine/Growth Factor 38-plex milliplex kit (Merck, PRCYTA-40K-PX38) for sCD137, CD40L, sFASL, G-CSF, GM-CSF, Granzyme A, Granzyme B, IFN-α2, IFN-γ, IL-1β, IL-1RA, IL-2, IL-4, IL-5, IL-6, IL-7, IL-8, IL-10, IL-12 (p70), IL-15, IL-17A, IL-18, IL-21, IL-22, IL-23, IL-33, IP-10, I-TAC, MCP-1, MIG, MIP-1α, MIP-1β, MIP-3α, Perforin, RANTES, TGFα, TNF-α, VEGF-A and a Bioplex 200 analyzer (Bio-Rad) according to manufacturer’s instructions. NHP-anti-virus response Legendplex kits (Biolegend) were also used. Dosages were performed following manufacturer recommendations. Final concentrations were obtained using manufacturer’ software.

### SIV viral load quantification

Plasma from whole blood samples was obtained by centrifugation at 1500g for 10 min. SIV RNA was isolated from plasma using a Nucleospin 96 Virus Core kit (Macherey-Nagel) or a QIAamp UltraSens Virus kit (Qiagen), according to manufacturer specifications. Quantitative RT-PCR using primers and probe towards the *gag* region of SIV genomic RNA were used to quantify plasma viral load, as previously described ^31^.

### Statistical analysis and data transformation

Cells count was determined using frequency among CD45^+^ cells, from whole blood cells, reported to complete blood count of leukocytes.

Normalizations of anti-LILRB1/B2 staining during competition assays were performed by using the formula:

All statistical analyses were performed using Prism 10 (GraphPad Software). Nonparametric Mann-Whitney U test, or a paired nonparametric Wilcoxon signed-rank test, were performed between unpaired or paired data, respectively. Friedman test was used to analyze longitudinal data. Comparisons of data between groups, time points, or pooled sample analyses used a Kruskall Wallis test and post-hoc tests.

## Acknowledgments

This work was supported by the Programme Investissements d’Avenir (PIA), managed by the ANR under reference ANR-11-INBS-0008, funding the Infectious Disease Models and Innovative Therapies (IDMIT, Fontenay-aux-Roses, France) infrastructure, and ANR-10-EQPX-02-01, funding the FlowCyTech facility (IDMIT, Fontenay-aux-Roses, France); the French National Agency for AIDS, Hepatitis and Emerging Infectious Disease (ANRS-MIE; grant number: ECTZ159192); and Sidaction (grant number: 21-1-AEQ-12963). The program is part of the “Institut Hospitalo-Universitaire, Comprehensive SEPSIS center”, funded by Agence Nationale de la Recherche (ANR, France) under reference IHU3 - ANR-23-IAHU-004. We thank the staff of the animal facility of IDMIT, particularly B. Delache, S. Langlois, J.M. Robert, Q. Sconosciuti, and N. Dhooge. We also thank O. Haigh for critical reading of the manuscript and W. Gros, M. Leonec and M. Van Tilbeurgh for flow cytometry assistance and L. Bossevot, M. Leonec, L. Moenne-Loccoz, and J. Morin for the RT-qPCR and Luminex assays. We also thank Valerie Monceaux for technical assistance.

---

**Supplementary table 1:** Antibodies used for phenotypic panels. Panels description: All = used for PK and SIV studies; Blood = Used only in blood panel; CA = used for cross-reactivity/cellular distribution assay; LN = Used for lymph node staining; PK = used for pharmacokinetic studies; SIV = used during SIV study.

| Target | clone | fluorochrome | manufacturer | reference | panel |
| --- | --- | --- | --- | --- | --- |
| CADM1 | PE | 3E1 | MBL | CM004-5 | All + CA |
| CD123 | PerCP-Cy5.5 | 7G3 | BD | 558714 | All + CA |
| CD14 | BV711 | M5E2 | Biologend | 301838 | All + CA |
| CD159a | PE-Vio770 | REA110 | Miltenyi | 130-113-567 | Blood; SIV |
| CD16 | BV650 | 3G8 | Biologend | 302042 | All + CA |
| CD163 | PE-Cy7 | GHI/61 | Biologend | 333614 | LN |
| CD1c | AF700 | L161 | Biologend | 331530 | All + CA |
| CD20 | PacificBlue | 2H7 | Biologend | 302328 | All |
| CD3 | Pacific Blue | SP34-2 | BD | 558124 | All |
| CD40 | FITC | 5c3 | Biologend | 334306 | SIV |
| CD45 | V500 | D058-1283 | BD | 561489 | All + CA |
| CD45RA | PE-Cy7 | 5H9 | BD | 561216 | SIV |
| CD66a,b,c,e | VioBlue | TET2 | Miltenyi | 130-119-851 | Blood; SIV |
| CD80 | BUV395 | L307.4 | BD | 565210 | SIV |
| CD86 | BUV737 | BU63 | BD | 748376 | SIV |
| CD95 | BV785 | DX2 | Biologend | 305646 | SIV |
| HLA-DR | BUV661 | L243 | BD | 753687 | SIV |
| HLA-DR | PE-Cy7 | L243 | BD | 335795 | PK + CA |
| HLA-I | BV605 | W6/32 | Biologend | 311432 | SIV |
| Live/Dead | BlueVid |  | Invitrogen | L34962 | All + CA |
| mLILRB1/B2 | AF647 | ch20G10;<br>mac20G10 | Fusion<br>antibodies;<br>biotem | homemade | All + CA |
| PD-L1 | BV421 | 29E.2A3 | BD | 568320 | SIV |
| Siglec1 | PEdazzle594 | 7-239 | Biologend | 346016 | SIV |

**Figure S01.**
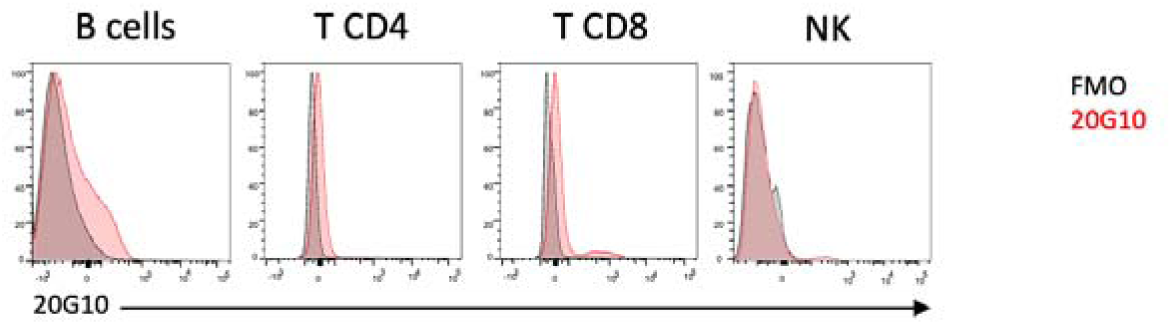
20G10 mAb staining distribution on blood lymphoid cells from cynomolgus macaque. Distribution of 20G10 mAb staining was assessed on PBMCs from cynomolgus macaque using flow cytometry.

**Figure S02.**
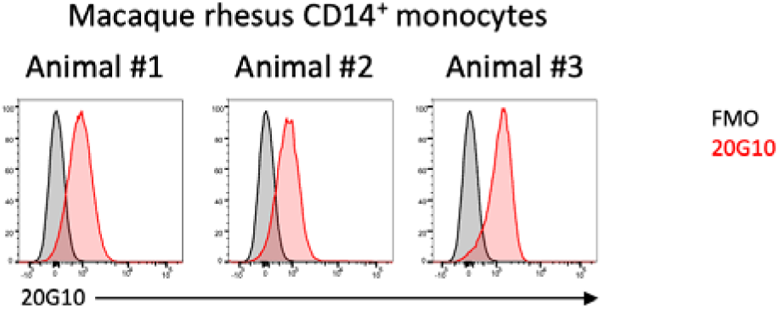
Cross-reactivity of 20G10 mAb with rhesus macaque. Cross-reactivity of 20G10 mAb with rhesus macaque monocytes was assessed by flow cytometry. PBMCs from rhesus macaque were used for staining of CD14^+^ monocytes (n=3 animals).

**Figure S03.**
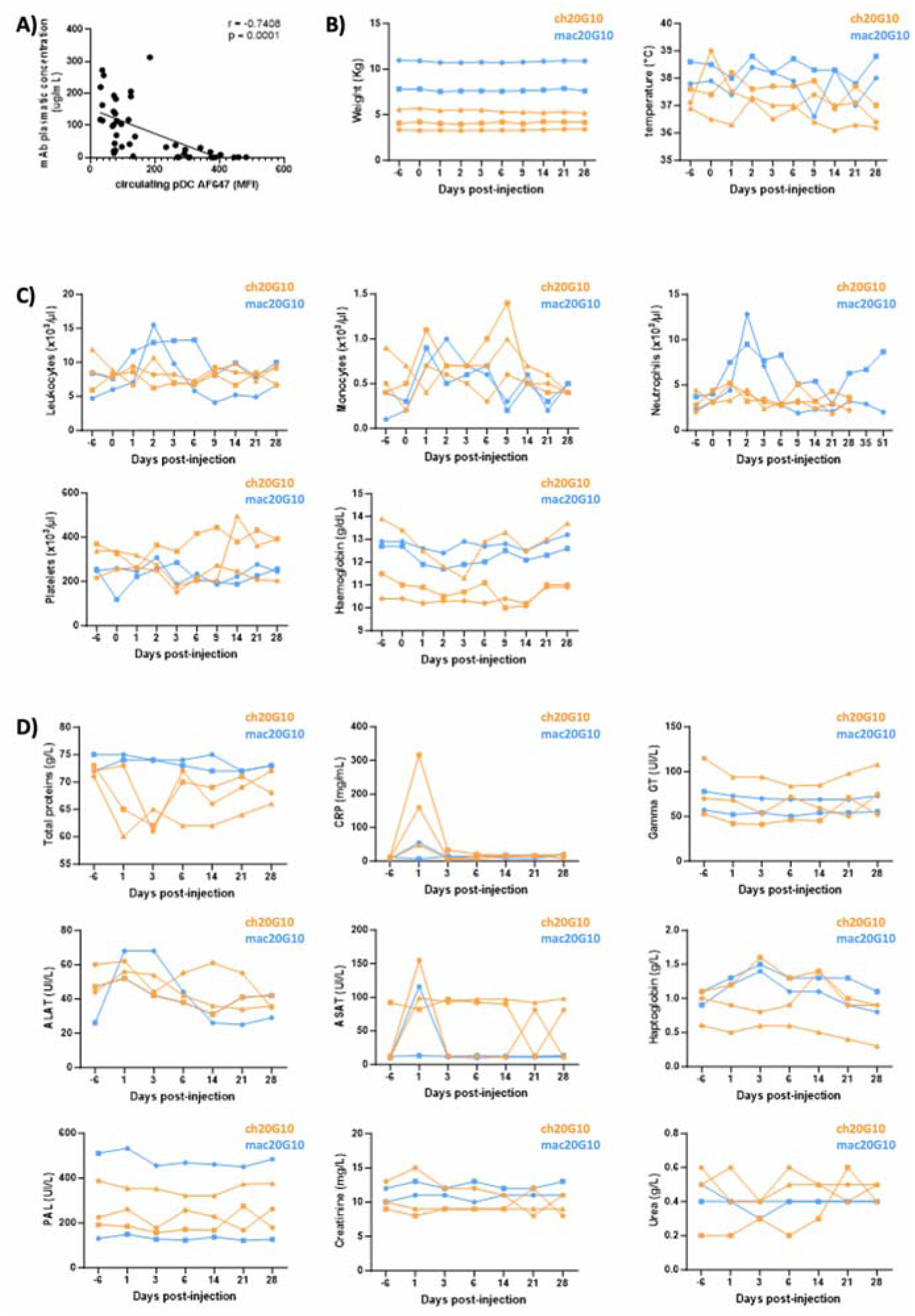
**Longitudinal analysis of pharmacodynamic parameters following ch20G10 or mac20G10 administration in cynomolgus macaques.** A) Correlation between plasma persistence of ch20G10 and mac20G10 and competition assay performed on circulating pDCs. Non-parametric spearman correlation test was used. B-D) Monitoring of animal weight and temperature (B), complete blood count (C) and biochemical parameters (D) after administration of ch20G10 (orange, n=3) or mac20G10 (blue, n=2).

**Figure S04.**
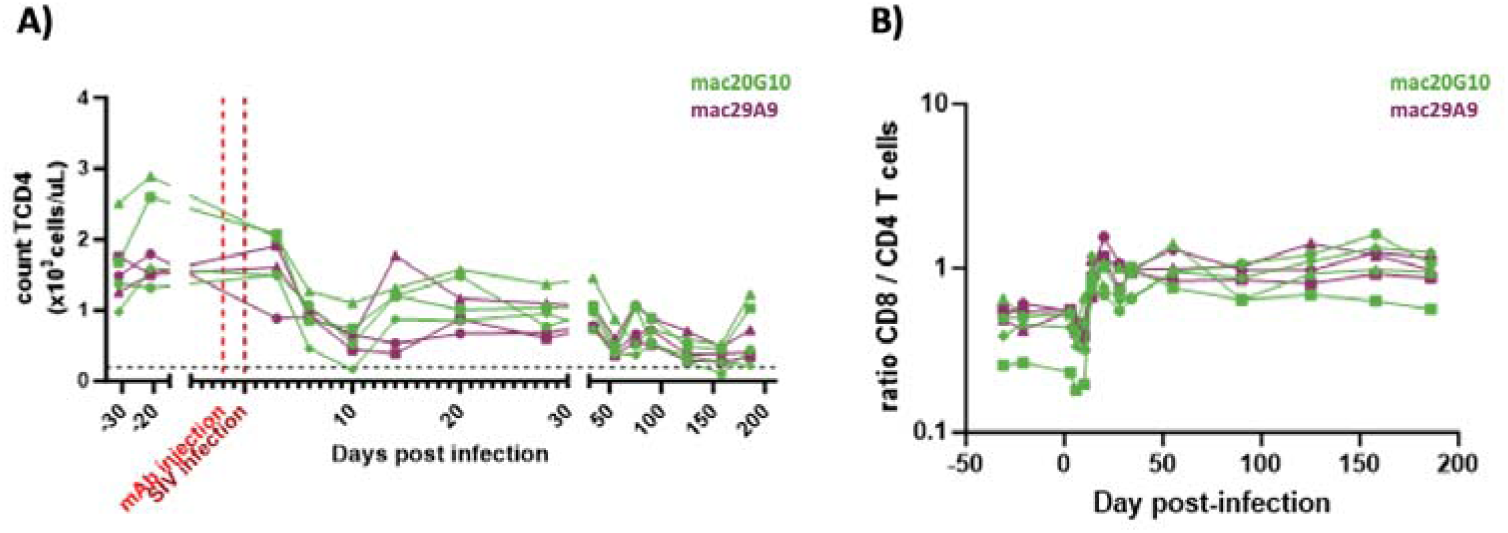
**Follow-up of CD4^+^ T cells in SIV-infected cynomolgus macaques.** A) Monitoring of CD4^+^ T cells absolute count per µL of blood determined by flow cytometry and complete blood count. B) Ratio of CD8^+^ T cell/CD4^+^ T cell in blood. mac20G10 treated group is in green (n=4) and mac29A9 isotype control group is in purple (n=3).

**Figure S05.**
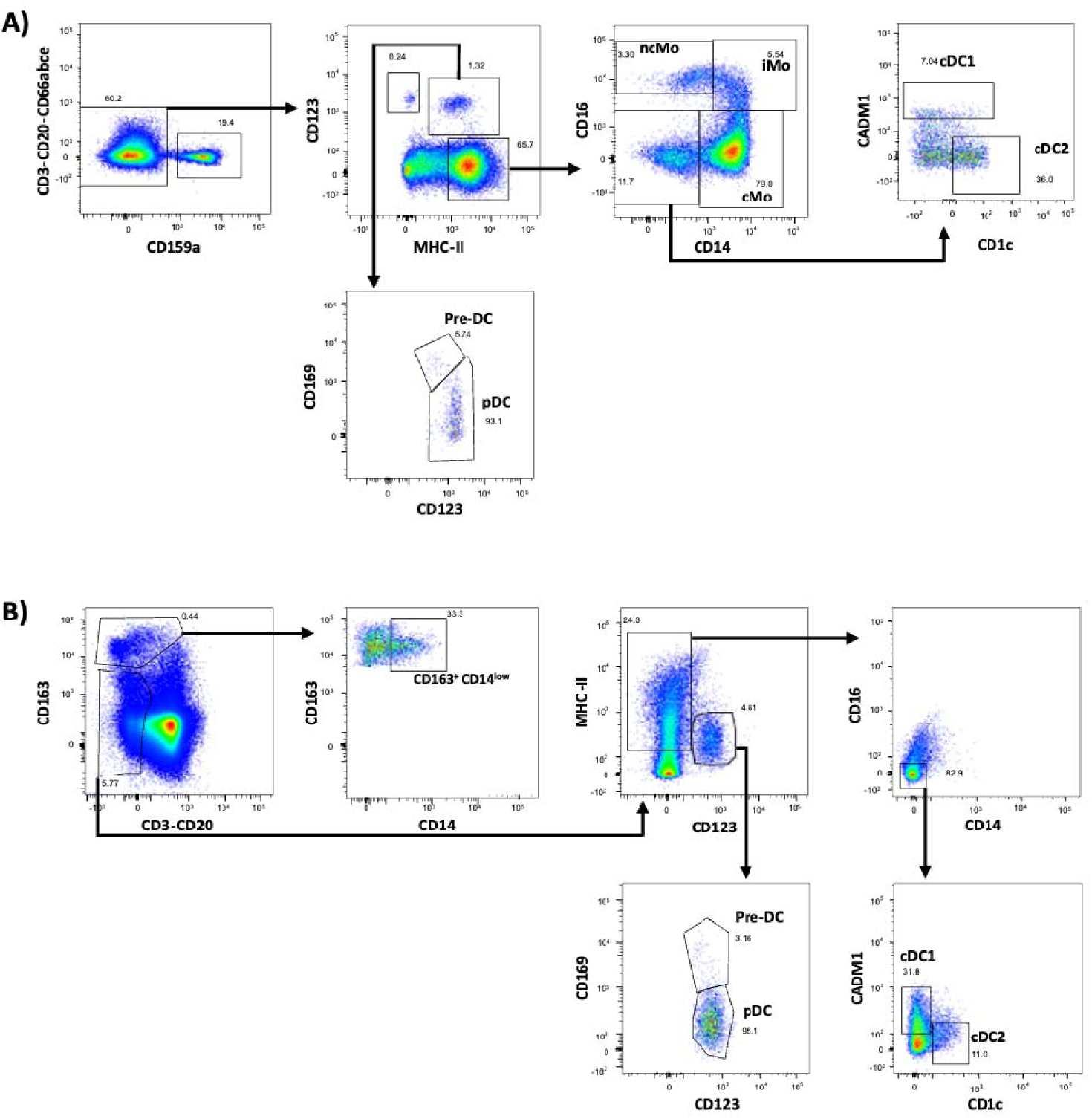
**Flow cytometry strategy used to follow myeloid subsets in blood and lymph nodes of cynomolgus macaques.** A) Flow cytometry strategy used to isolate cDC1, cDC2, pDC and monocytes subsets in blood. cMo = classical monocytes, iMo = inflammatory monocytes and ncMo = non-classical monocytes. B) Flow cytometry strategy used to isolate cDC1, cDC2, pDC and macrophages in lymph nodes.

**Figure S06.**
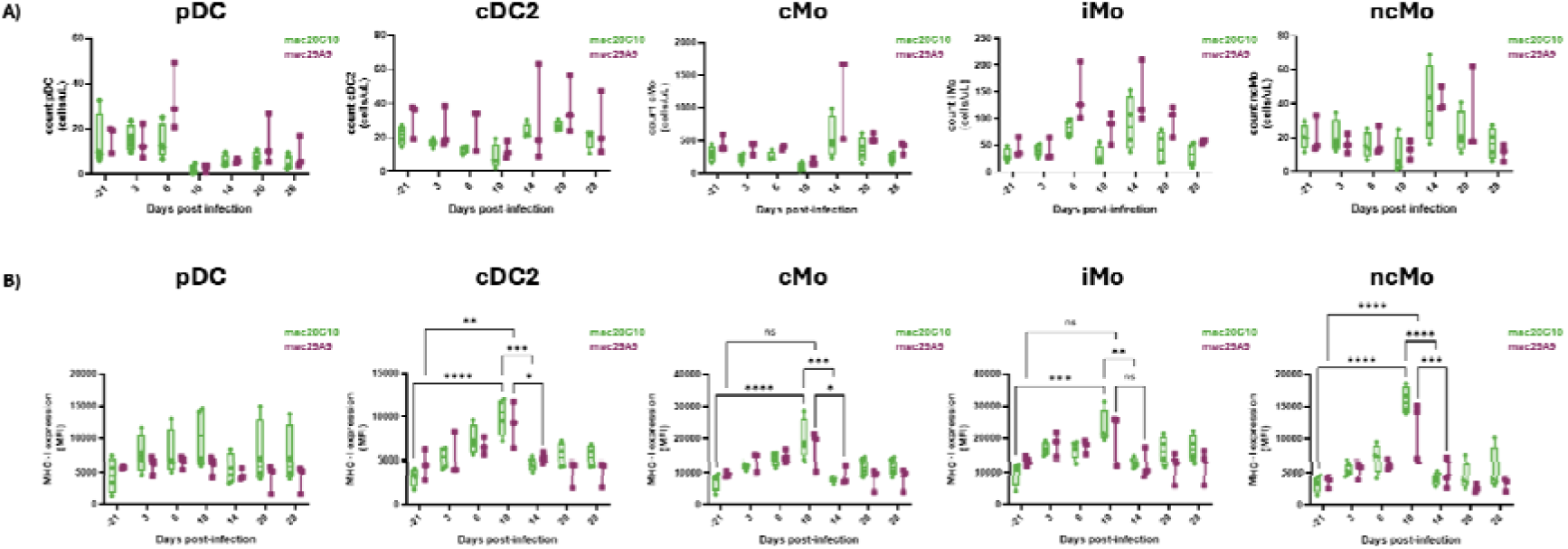
**Follow-up of pDC, cDC2 and monocytes count and their expression of MHC-I expression in early SIV infection of macaques treated with mac20G10 or mac29A9.** A) Count of pDC, cDC2 and monocytes subsets during SIV infection. cMo = classical monocytes, iMo = inflammatory monocytes and ncMo = non-classical monocytes. B) Evolution of MHC-I expression on pDC, cDC2 and monocytes sub-populations measured by flow cytometry. Groups and time points were compared using a Kruskal-Wallis test, followed by post-hoc analysis; * = p<0.05, ** = p<0.01, *** p<0.001, **** p<0.0001. mac20G10 treated group is in green (n=4) and control isotype group is in purple (n=3).

**Figure S07.**
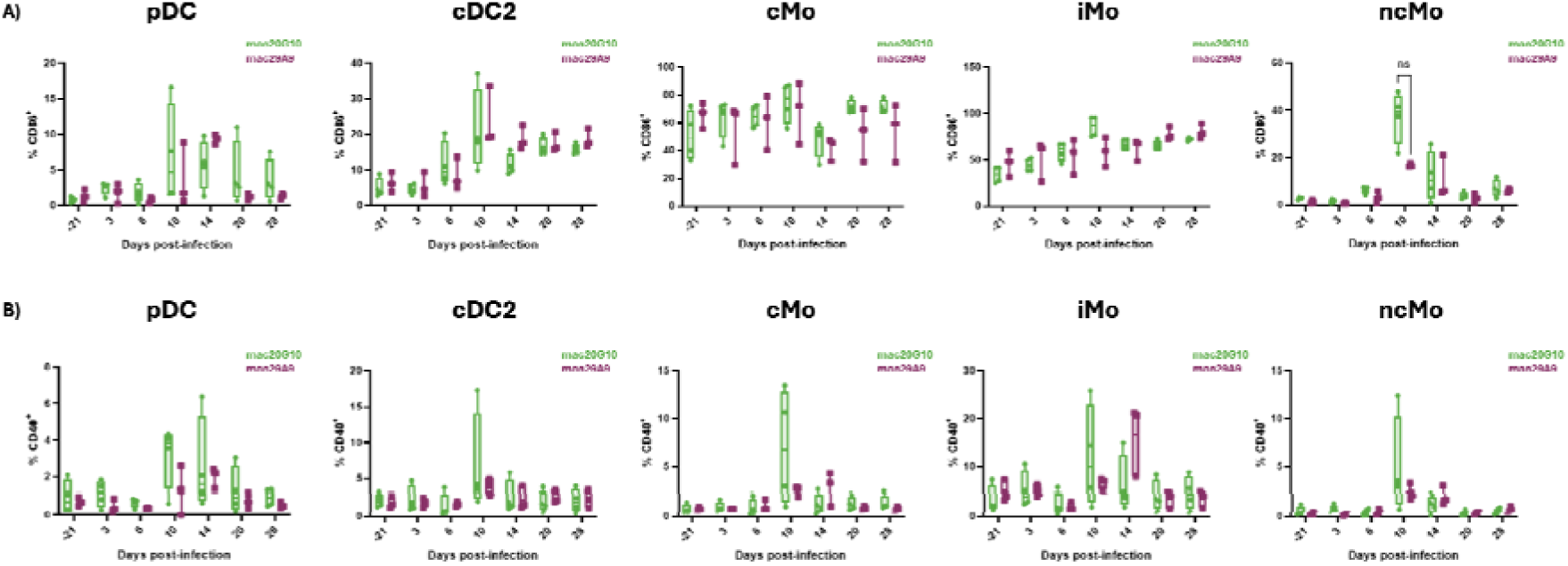
**Follow-up of the frequency of CD86^+^ and CD40^+^ myeloid subsets from SIV-infected cynomolgus macaques treated with mac20G10 or mac29A9.** A) Follow up of CD86^+^ pDC, cDC2 and monocyte sub-populations frequencies during acute phase of infection. cMo = classical monocytes, iMo = inflammatory monocytes and ncMo = non-classical monocytes. Groups and time points were compared using a Kruskal-Wallis test, followed by post-hoc analysis. B) Follow up of the CD40^+^ pDC, cDC2 and monocytes subpopulations frequencies during acute phase of infection. Groups and time points were compared using a Kruskal-Wallis test, followed by post-hoc analysis. mac20G10 treated group is in green (n=4) and control isotype group is in purple (n=3).

**Figure S08.**
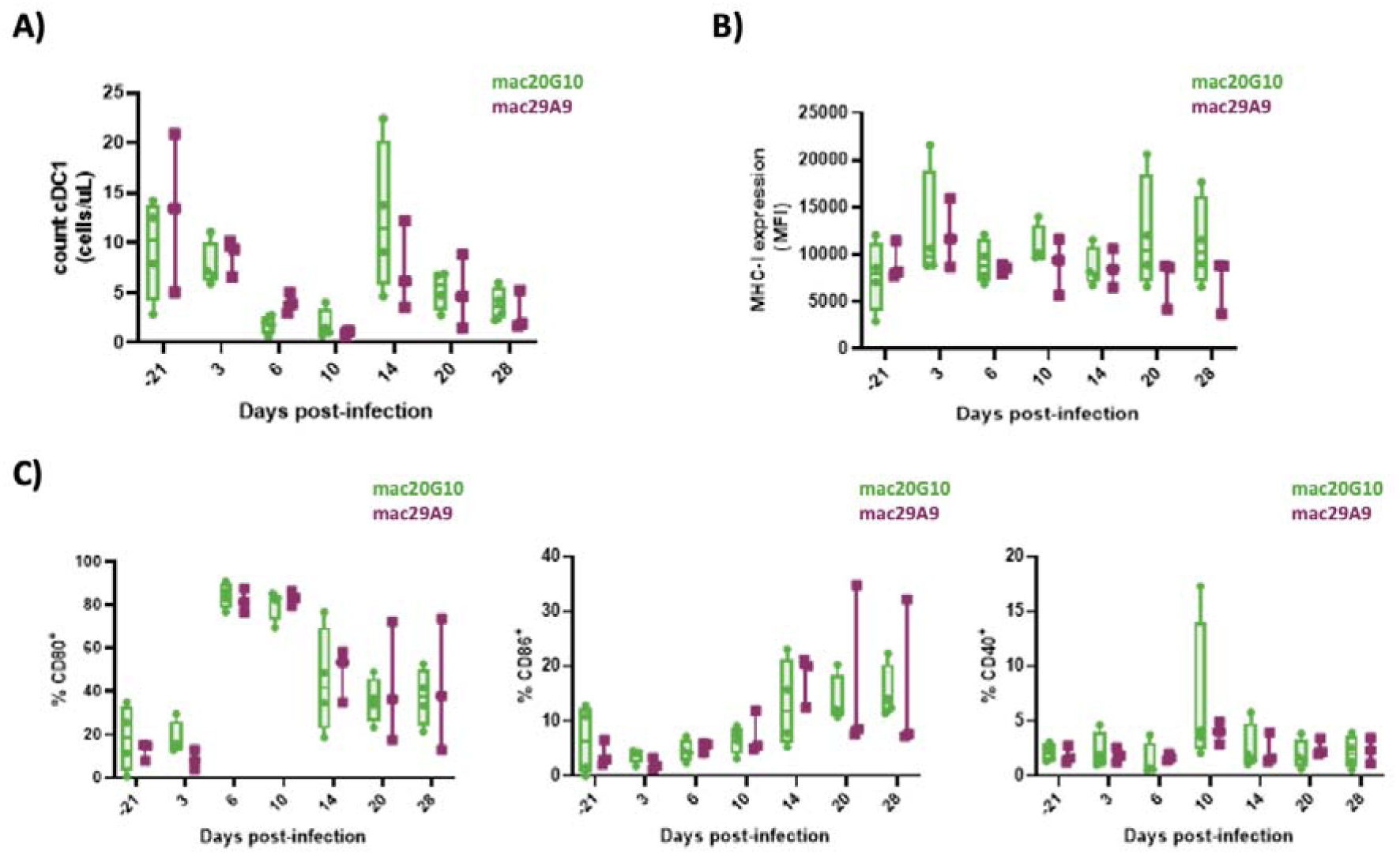
**Follow-up of cDC1 count and phenotype from SIV-infected cynomolgus macaques treated with mac20G10 or mac29A9.** A) Count of cDC1 during SIV infection. B) Evolution of MHC-I expression on cDC1, measured by flow cytometry. C) Follow up of CD80^+^, CD86^+^ or CD40^+^ cDC1 frequencies. Groups and time points were compared using a Kruskal-Wallis test, followed by post-hoc analysis; * = p<0.05, ** = p<0.01, *** p<0.001, **** p<0.0001. mac20G10 treated group is in green (n=4) and control isotype group is in purple (n=3).

**Figure S09.**
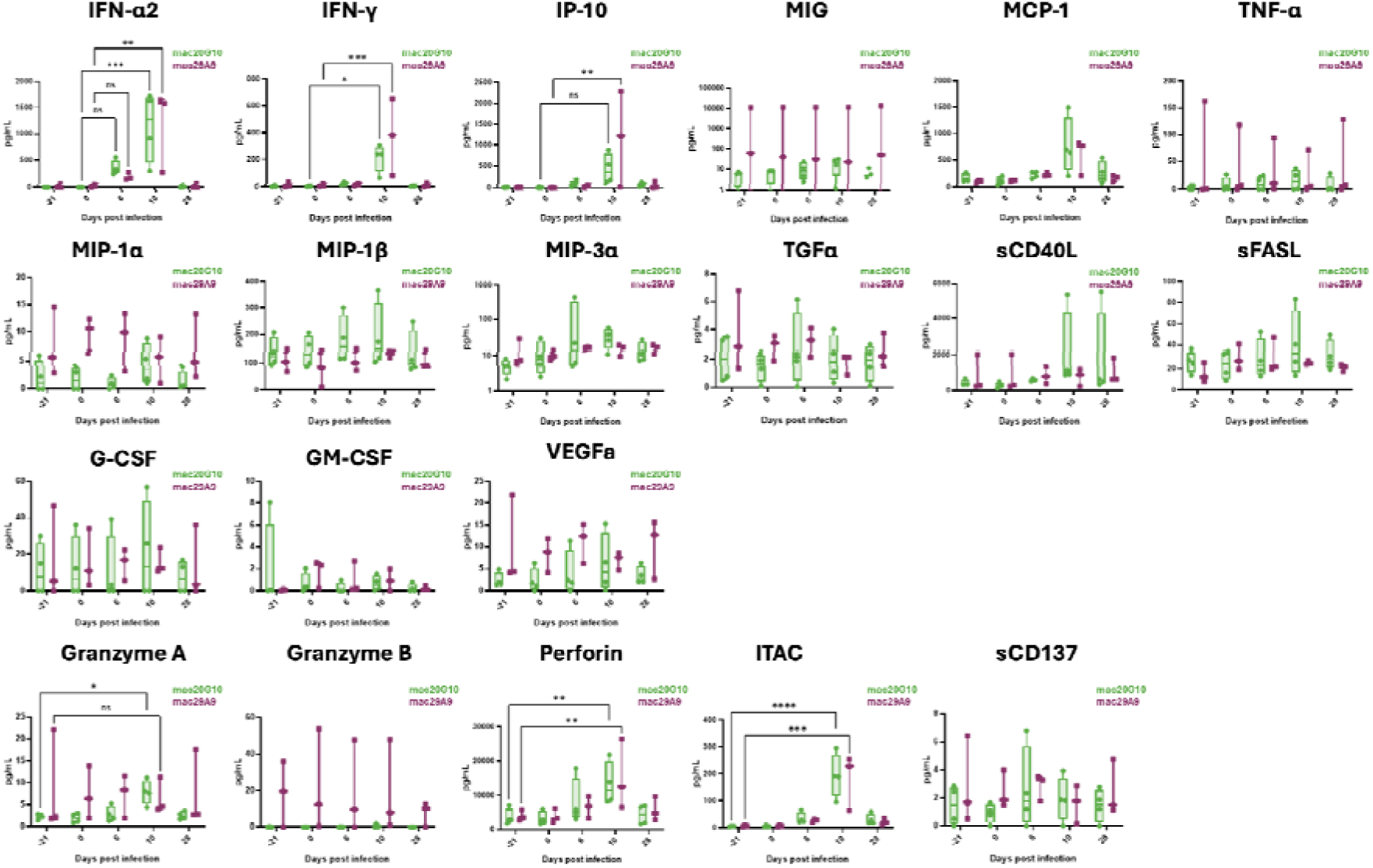
**Follow-up of plasma cytokines in early SIV infection from cynomolgus macaques treated with mac20G10 or mac29A9.** Concentrations of IFN-α2, IFN-γ, IP-10, MIG, MIP-1α, MIP-1β, MIP-3α, MCP-1, TGFα, TNF-α, soluble CD40 ligand (sCD40L), soluble FAS ligand (sFASL), G-CSF, GM-CSF, VEGF-a, Granzyme A, Granzyme B, Perforin, ITAC and CD137 were measured using Luminex. Groups and time points were compared using a Kruskal-Wallis test, followed by post-hoc analysis; * = p<0.05, ** = p<0.01, *** p<0.001, **** p<0.0001. mac20G10 treated group is in green (n=4) and control isotype group is in purple (n=3).

**Figure S10.**
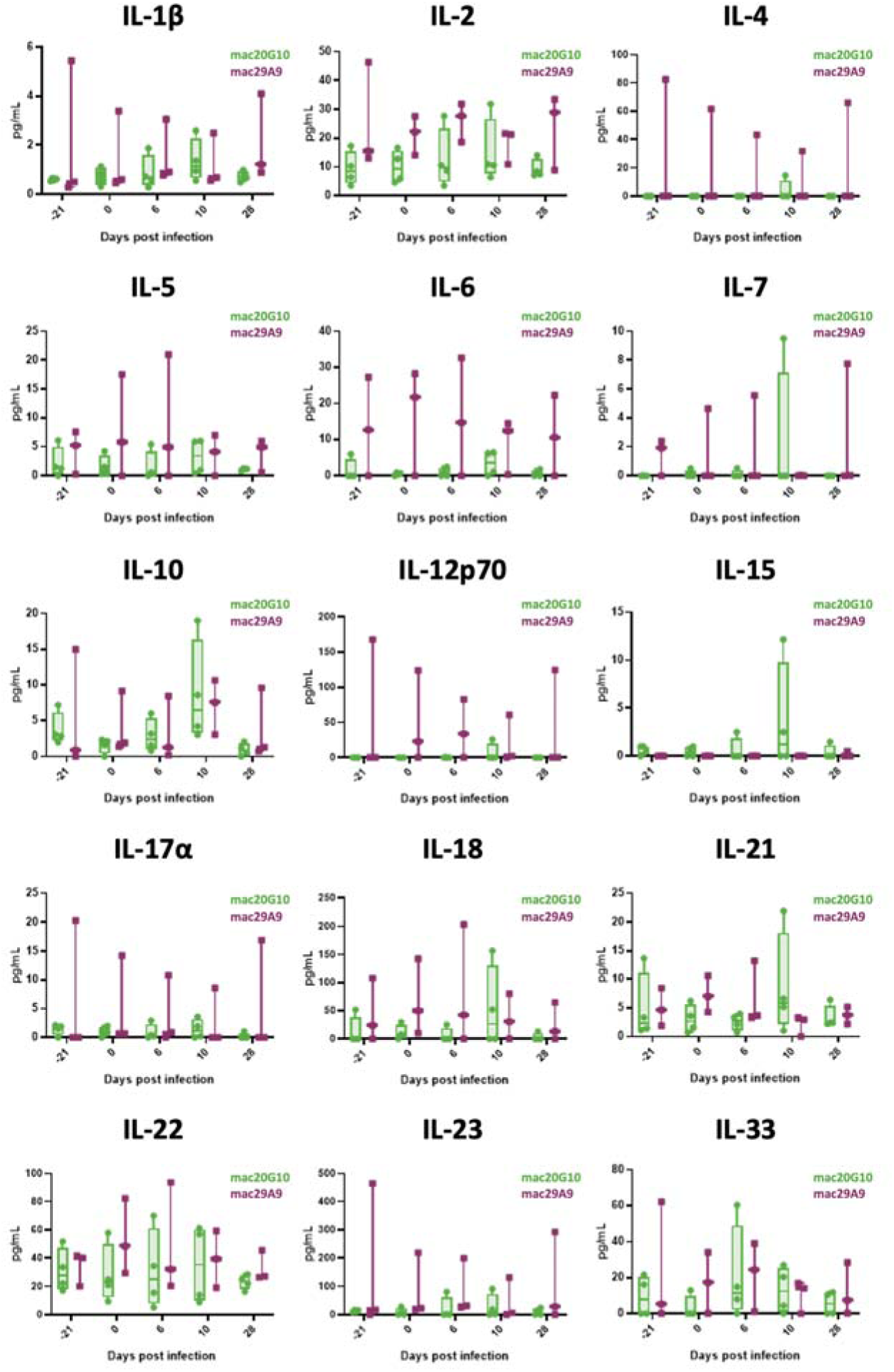
**Follow-up of plasma interleukins in early SIV infection of cynomolgus macaques treated with mac20G10 or mac29A9.** Concentrations of IL-1β, IL-2, IL-4, IL-5, IL-6, IL-7, IL-10, IL-12p70, IL-15, IL-17α, IL-18, IL-21, IL-22, IL-23, IL-33 were measured by Luminex. Groups and time points were compared using a Kruskal-Wallis test, followed by post-hoc analysis. mac20G10 treated group is in green (n=4) and control isotype group is in purple (n=3).

**Figure S11.**
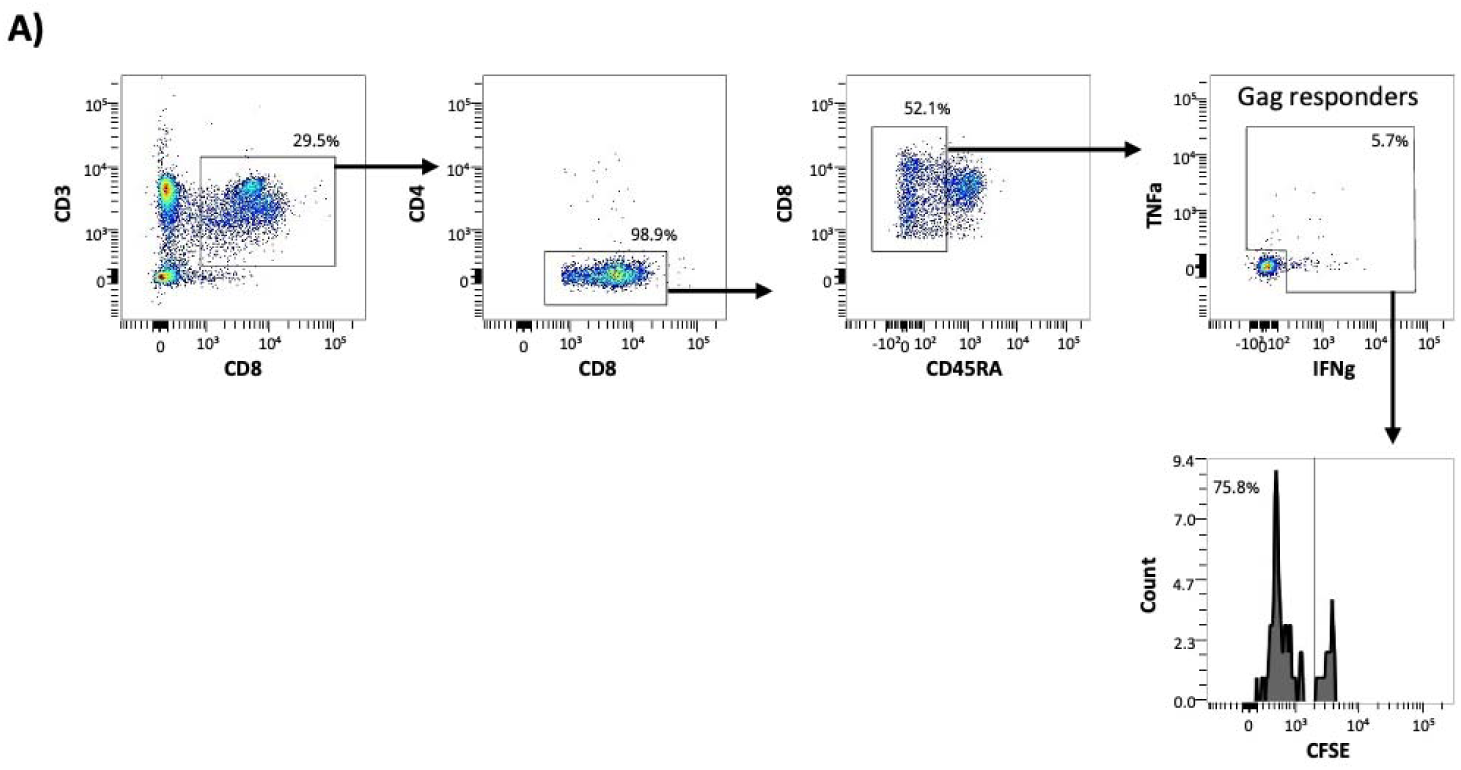
**Flow cytometry gating strategy used for the characterization of SIV^gag^ specific memory CD8^+^ T cells after ex vivo stimulations.**

## Notes

### Competing Interest Statement

The authors have declared no competing interest.

